# Human cardiac fibrosis-on-a-chip model recapitulates disease hallmarks and can serve as a platform for drug screening

**DOI:** 10.1101/632406

**Authors:** Olya Mastikhina, Byeong-Ui Moon, Kenneth Williams, Rupal Hatkar, Dakota Gustafson, Xuetao Sun, Margaret Koo, Alan Y.L. Lam, Yu Sun, Jason E. Fish, Edmond W.K. Young, Sara S. Nunes

## Abstract

While interstitial fibrosis plays a significant role in heart failure, our understanding of disease progression in humans is limited. To address this limitation, we have engineered a cardiac-fibrosis-on-a-chip model consisting of a microfabricated device with live force measurement capabilities using co-cultured human cardiac fibroblasts and pluripotent stem cell-derived cardiomyocytes. Transforming growth factor-β was used as a trigger for fibrosis. Here, we have reproduced the classic hallmarks of fibrosis-induced heart failure including high collagen deposition, increased tissue stiffness, BNP secretion, and passive tension. Force of contraction was significantly decreased in fibrotic tissues that displayed a transcriptomic signature consistent with human cardiac fibrosis/heart failure. Treatment with an anti-fibrotic drug decreased tissue stiffness and BNP secretion, with corresponding changes in the transcriptomic signature. This model represents an accessible approach to study human heart failure *in vitro*, and allows for testing anti-fibrotic drugs while facilitating the real-time assessment of cardiomyocyte function.

## Introduction

Heart failure is the end-stage clinical manifestation of multiple forms of cardiovascular diseases. It is characterized by cardiac remodeling and reduced ventricular compliance as a consequence of interstitial fibrosis (Bloom et al., 2017). Heart damage caused by various insults (i.e. hypoxia during myocardial infarction, MI) triggers complex wound healing mechanisms to restore homeostasis and while collagen deposition is a normal and essential component of wound healing, it can evolve into a progressively irreversible fibrotic response. In the setting of cardiac fibrosis, fibroblast activation leads to excessive deposition of extracellular matrix components such as collagen and fibronectin, and increases tissue stiffness, which further activates cardiac fibroblasts and results in a positive feedback loop of activation and increased stiffness (Travers et al., 2016). This can compromise heart function and increase the risk of arrhythmia, ultimately leading to heart failure.

Although a number of factors have been implicated in orchestrating the fibrotic response, tissue fibrosis is dominated by a central mediator: transforming growth factor-β (TGF-β (Meng et al., 2016)). Sustained TGF-β secretion leads to a continuous cycle of growth factor signaling and dysregulated matrix turnover. Despite abundant evidence from animal models of fibrosis and samples from human biopsies supporting a central role for TGF-β in tissue fibrosis, modulating TGF-β function to attenuate or reverse fibrosis is replete with challenges. For example, attempts to block TGF-β signaling using antagonists to the type I activin receptor-like kinase 5 (ALK5) have failed to reach clinical studies, owing mainly to safety concerns (Anderton et al., 2011). Therefore, it is necessary to improve our understanding of disease progression to uncover alternative targets and develop strategies to treat cardiac fibrosis.

Despite advances in our knowledge of fibroblast activation and fibrosis due to research performed in animal models and *in vitro* studies on fibroblast activation, our understanding of disease progression in humans is limited. Moreover, animal models often fail to faithfully mimic human responses, while reductionist approaches at the cell level are most commonly used in conventional 2D in vitro platforms that fail to recapitulate higher dimensionality as well as higher order intercellular interactions. This motivates the need for a human biomimetic *in vitro* platform to investigate the progression of fibrotic remodeling in three dimensions (3D) that allows for real-time monitoring of functional outcomes. Such near-physiological models enable the study of a range of biological processes such as cell-cell interactions and signaling that take place in the 3D environment, as well as tissue responses to drugs (Ingber, 2016). They also allow for controlled studies of organ-level aspects of human physiology and disease, and have been successfully used to uncover new potential therapeutic targets (Huh et al., 2012).

We and others have generated “healthy” and disease models of human cardiac tissues *in vitro* from hiPSC-CMs, including long-QT and Barth syndromes (Itzhaki et al., 2011; MacQueen et al., 2018; Moretti et al., 2010; Nunes et al., 2013; Ronaldson-Bouchard and Vunjak-Novakovic, 2018; Tiburcy et al., 2017; Wang et al., 2014; Zhao et al., 2019). However, no human models of fibrosis-induced heart failure have been generated. Instead, models described to date have been of animal origin (Desroches et al., 2012; Sadeghi et al., 2017) requiring interspecies extrapolation, or lack assessment of key functional parameters such as force of contraction and anti-fibrotic drug testing (Lee et al., 2019)

Here, we describe for the first time and thoroughly characterize the development of a 3D model of human cardiac fibrosis (hCF-on-a-chip) using human cardiac fibroblasts together with human induced pluripotent stem cell-derived cardiomyocytes (hiPSC-CMs) in a platform with live imaging capabilities and real-time assessment of contractile function. The resulting hCF-on-a-chip technology displayed the hallmark characteristics of cardiac fibrosis and associated heart failure including impaired functional capacity, increased collagen deposition and tissue stiffness, increased brain natriuretic peptide (BNP) secretion and a ‘fibrotic’ transcriptomic signature, including changes in key microRNAs secreted in extracellular vesicles and implicated in human cardiac fibrosis. Moreover, we show that standard-of-care drugs losartan and carvediolol significantly decreased BNP secretion in the 3D hCF-on-a-chip tissues, but not in 2D monolayers. In addition, pirfenidone, a commercially available drug for treatment of idiopathic pulmonary fibrosis (Noble et al., 2011) (IPF), significantly reduced tissue stiffness and BNP secretion, as well as changed the microRNA signature in tissues that had already displayed significant functional capacity loss. If performed for longer time, pirfenidone treatment also led to improvement is passive tissue tension.

## Results

### Human cardiac fibrosis-on-a-chip (hCF) design, fabrication and characterization

To recapitulate cardiac fibrosis *in vitro* and concurrently facilitate force of contraction measurements, we developed and used an accessible, two-material microwell chip consisting of a cell culture compartment and two parallel flexible horizontal rods at each end of the culture compartment (**Fig. 1a**). The device was fabricated in poly(methylmethacrylate) (PMMA or acrylic) or polystyrene using micromilling methods in thermoplastics (Guckenberger et al., 2015). The two flexible rods were made from poly(dimethylsiloxane) (PDMS), and fabricated by withdrawing mixed PDMS prepolymer (PDMS base and curing agent) into a 27-gauge syringe needle (outer diameter ∼400 μm, inner diameter ∼200 μm), curing the PDMS inside the needle, and pulling the cured PDMS out of the syringe needle to produce cylindrical rods (**Supplemental Fig. 1a**). The platform is open-well and pipette-accessible as in (Conant et al., 2017). Real-time monitoring of rod deflection is facilitated by the horizontal orientation of the rods, enabling calculation of both passive tissue tension (relaxation) and active force (contraction). The device consists of three microwell systems each containing a pair of the PDMS rods. The rods were secured by interference fits into microscale notches in the PMMA/polystyrene layer (**Supplemental Fig. 1a**).

**Figure 1.**
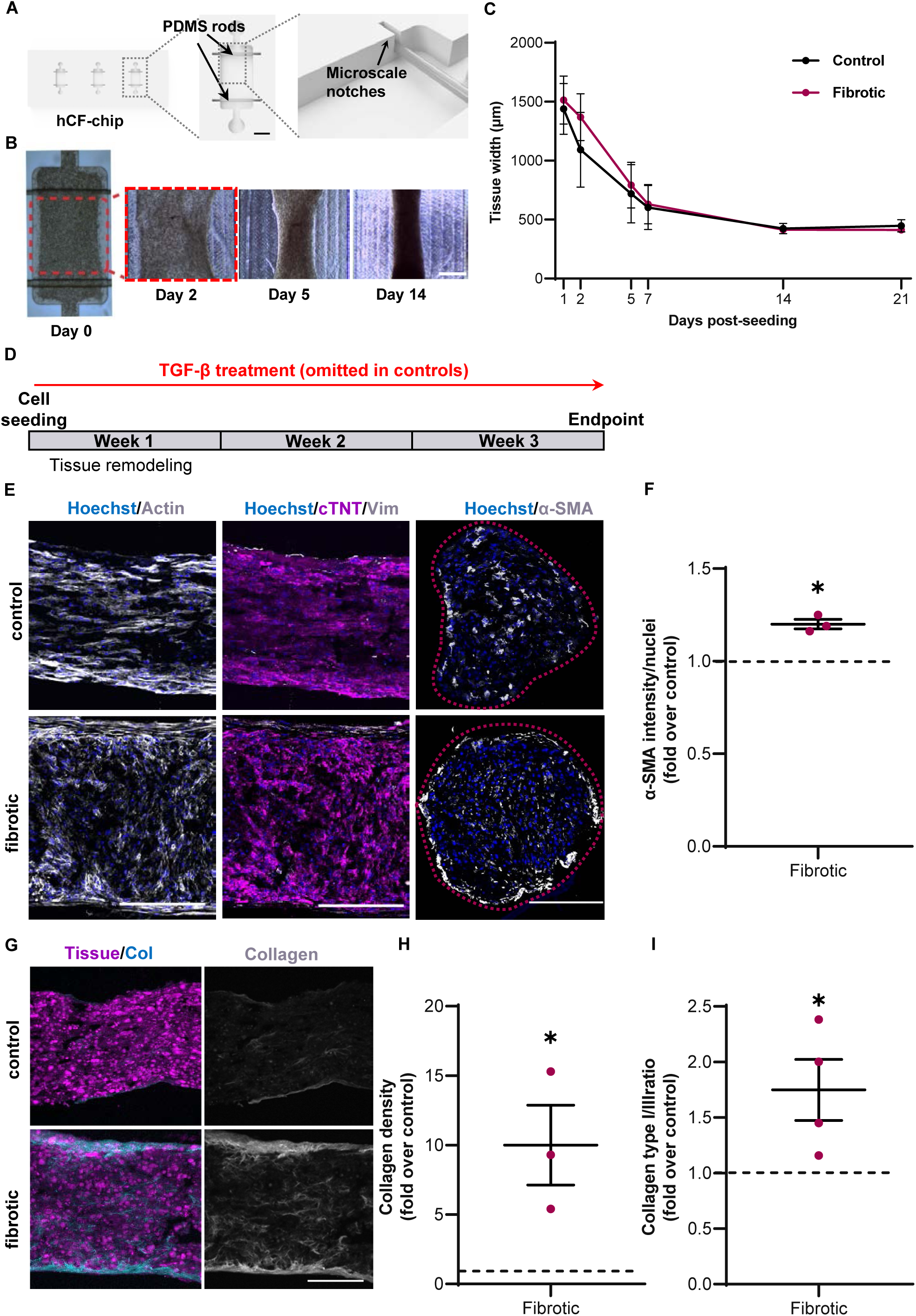
Development and characterization of the human cardiac fibrosis-on-a-chip model. **a**, Render of the hCF-chip containing 3 microwells (left image). Top view image shows a well with two elastomeric rods (arrows) suspended on each end (middle image). Scale bar, 1 mm. Schematic of the microscale notches present on each end of the well, used to support the elastomeric rod in place (right image). **b**, Representative images for control tissues at days 0, 2, 5, and 14 post-seeding. Scale bar: 500 μm. **c**, Central tissue width variation over time of control (black) and fibrotic tissues (red). **d**, Schematic of the conditions and timeline of tissue treatments. hiPSC-CMs were seeded with untreated (control) or TGF-β1-treated (fibrotic) human cardiac fibroblasts. Microtissues were allowed to self-assemble during the first week in culture and were assessed throughout time up to 3 weeks post-seeding (endpoint). **e**, Cardiac tissue architecture and cell distribution was assessed by staining longitudinal tissue sections for nuclei (Hoechst, blue) actin (phalloidin, white), cardiac troponin T (magenta) and vimentin (grey). Tissue transverse sections were stained for nuclei (Hoechst, blue) and α-SMA (grey). Red dashed line represents tissue outline. Scale bars: 200 μm. **f**, Quantification of α-SMA staining at endpoint shows fibrotic tissues had significantly higher α-SMA expression vs control (dashed line) (average ± SEM, p = 0.014, N = 3). **g**, Representative images of control and fibrotic tissues assessed by second harmonic generation. Collagen (Col), cyan (left) and grey (right), and cells - magenta (auto-fluorescence). Scale bars: 250 μm. **h**, Quantification of second harmonic images for fibrotic tissues, area of collagen over tissue area. Fold vs. control (dashed line) (average ± SEM, p = 0.018, N = 3) **i**, Fibrotic tissues showed a significantly higher collagen type I/type III ratio than control as evaluated by collagen type I and type III immunohistochemistry quantification vs. control (dashed line) (average ± SEM, p = 0.049, N = 3).

A ratio of 3:1 hiPSC-CM to human cardiac fibroblasts was chosen to reflect the ratio of cardiomyocyte and non-cardiomyocyte cell populations in the heart (Pinto et al., 2016). Cells were seeded into the microwells in fibrin gels to alleviate the addition of exogenous collagen and to permit the accurate assessment of collagen deposited by the seeded cells. Gels with cells were allowed to self-assemble into 3D microtissues (**Fig. 1b**). Assessment of gel compaction demonstrated that tissues were fully compacted in the second week post-seeding (**Fig. 1b, c**), with 78% gel compaction. These microtissues (∼550 x 300 μm in size) beat synchronously and spontaneously, and were amenable to pacing by electrical field stimulation by placing the chip into customized electrical stimulation chambers (Nunes et al., 2013) (**Supplemental Fig. 1b, c**). Rod deflection can be quantified by microscopy and analysis of video recordings (**Supplemental Fig. 2a** and **Supplemental Videos 1-2**). This platform also permits the utilization of rods of different characteristics (i.e. rods made with different ratios of PDMS to curing agent), which allows for the tuning of the characteristics of the PDMS rod and to resolve different force ranges. For measuring force of contraction we used PDMS rods cured at a ratio of 1:30 of curing agent to PDMS base. Our setup resembles basic beam bending with distributed load at midspan and fixed constraints at the ends, but due to the elastomeric properties of the rod and large deflections, classic beam bending equations for linear elastic materials were not applicable. Instead, force of contraction was determined by measuring elastomeric rod deflection in recorded videos, and then converting the deflection to applied force (from a set of hanging weights) using standard multivariable polynomial curve-fit of deflection measurements (**Supplemental Fig. 2b**). The derived equation was validated with another set of PDMS rods and hanging weights (**Supplemental Fig. 2c**). Rod deflection at peak cardiomyocyte relaxation was used to assess passive force, while maximum deflection (during peak cardiomyocyte contraction) was used to quantify peak force of contraction by subtracting total force from passive tension (**Supplemental Fig. 2a, b**). Tissues beat spontaneously and responded to treatment with isoproterenol by increasing beating frequency (**Supplemental Fig. 3** and **Supplemental Videos 1, 2**).

### Recapitulation of tissue fibrosis

To replicate the contractile pathophysiology of cardiac fibrosis with human cells *in vitro*, we used human cardiac fibroblasts activated in plastic cell culture dishes with TGF-β1(Thannickal et al., 2003) co-seeded with hiPSC-CMs (this condition is henceforth referred to as ‘fibrotic’).

Untreated fibroblasts co-seeded with hiPSC-CMs were used as our control samples (‘control’). TGF-β1 was maintained in the medium in fibrotic samples and omitted in controls throughout the experiment course (**Fig. 1b**). After seeding in fibrin gels, cells were allowed to self-assemble (**Fig. 1b, d**, tissue remodeling stage). Areas with beating cells within the tissues were observed around day 2, and synchronous contraction was observed around day 5 post-seeding. Fibrotic tissues compacted at a similar rate compared to control (**Fig. 1c**). Several parameters were analyzed during weeks 2 and 3 post-seeding. Histological analysis was performed at the end of week 3 post-seeding (day 21, endpoint).

Human iPSC-CMs in the control tissues strongly expressed cardiac contractile proteins (actin) and cardiac troponin T, and showed alignment along the long axis of the tissues (**Fig. 1e**, control). Fibrotic tissues displayed a 20% increase in alpha smooth muscle actin (α-SMA) intensity relative to control (**Fig. 1f**) and had α-SMA-positive cells concentrated on the outer edge compared to a more homogeneous distribution of α-SMA-positive cells in controls at day 21 (**Fig. 1e**). Next, we assessed fibrillar collagen deposition, a hallmark of tissue fibrosis, by second harmonic generation imaging (Chen et al., 2012). There was an increase in interstitial collagen deposition, with fibrotic microtissues showing a 10-fold increase in fibrillar collagen content compared to control tissues (**Fig. 1g, h**). Moreover, consistent with clinical data (Bishop et al., 1990; Marijianowski et al., 1995), the ratio of collagen type I to III was significantly increased in fibrotic tissues compared to control (**Fig. 1i**). In agreement with increased collagen deposition and an increase in collagen type I/III ratio, average stiffness of live tissues assessed by atomic force microscopy was 2.3 times higher in fibrotic tissues when compared to controls (**Fig. 2a**)(Herum et al., 2017). Importantly, fibrotic tissues also had increased heterogeneity in local stiffness compared to control tissues, with stiffness ranging from 0.6 to 3.6 kPa in controls, and 0.6 to 9.2 kPa in fibrotic tissues (**Fig. 2b**). This is in line with the described heterogeneity of tissue fibrosis (Wells, 2013). Accordingly, passive tension (at peak cardiomyocyte relaxation, comparable to cardiac diastole) in fibrotic tissues was over 4-fold higher in fibrotic tissues compared to controls (**Fig. 2c**). Moreover, force of contraction in fibrotic tissues (active force, comparable to cardiac systole) was 3.1- and 5-fold lower in fibrotic tissues compared to control at days 14 and 21 post-seeding, respectively. (**Fig. 2d, Supplemental Videos 3, 4**). Excitation threshold, a measurement of cell electrical connectivity, was 1.75 times higher in fibrotic tissues compared to controls (**Fig. 2e**), indicating poorer cell connectivity. Moreover, levels of secreted BNP, an essential diagnostic marker and prognostic indicator in heart failure (Troughton et al., 2014), were significantly higher in fibrotic tissues compared to controls (**Fig. 2f**). No differences were observed in overall cell apoptosis or proliferation at day 21 post-seeding (**Supplemental Fig 4a, b, c**). The percentage of vimentin- and cTnT-positive cells, as well as proliferation and apoptosis was similar in fibrotic relative to control tissues at day 21 (**Supplemental Fig. 4a, d, e**), indicating that the 86% reduction in active force of contraction in fibrotic tissues relative to control at day 21 was not caused by a potential loss of cardiomyocytes.

**Figure 2.**
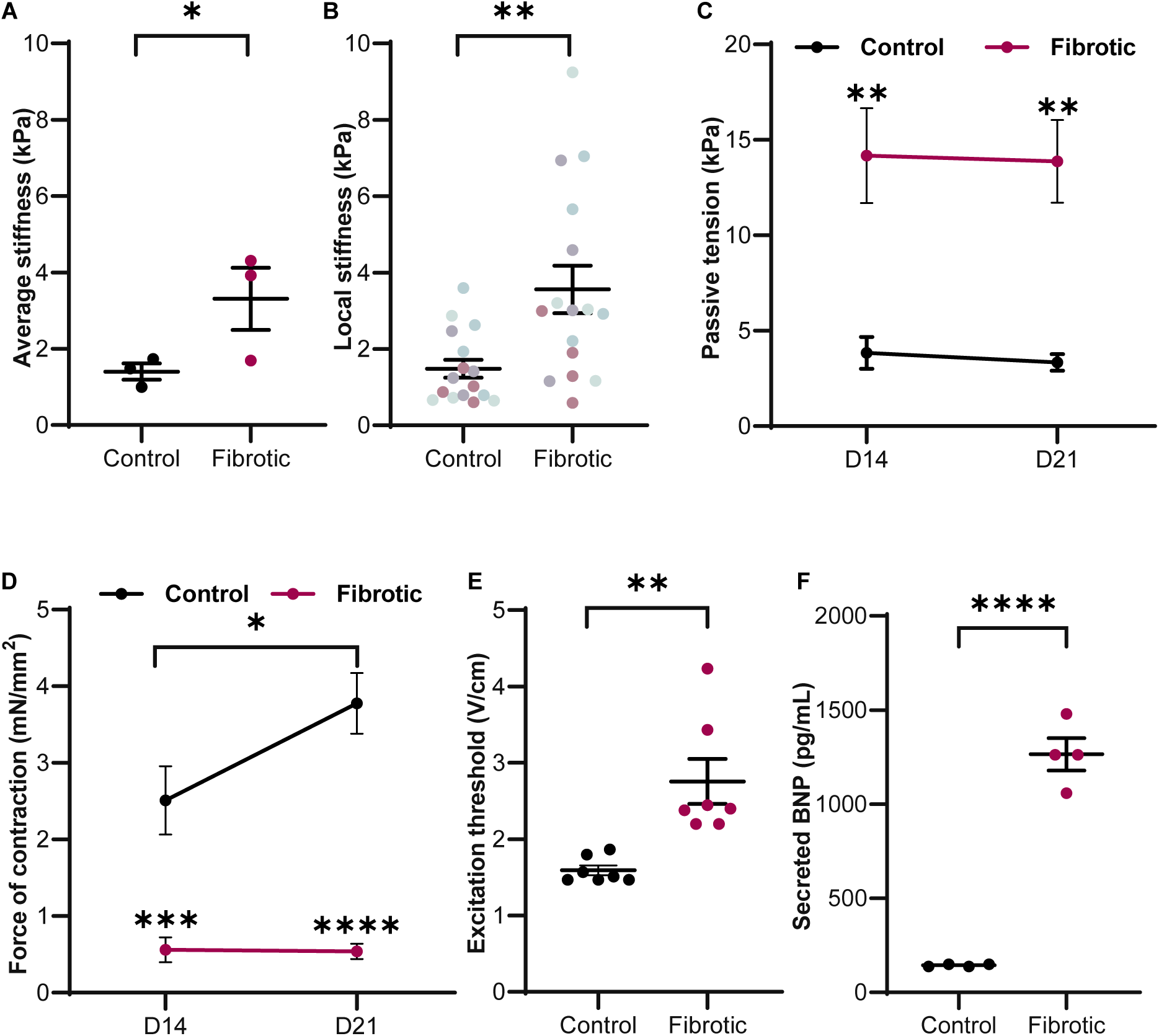
Characterization of the microtissues indicates fibrosis affects human cardiac tissue function. **a**, Average tissue stiffness assessed by atomic force microscopy at endpoint (average ± SEM, p = 0.029, N = 3). **b**, Individual stiffness values for each point probed shows increased heterogeneity in fibrotic tissues compared to controls (average ± SEM, p = 0.004, N = 3, n = 4 tissues total (each tissue represented in the same color), and 4 points/tissue). **c**, Passive tissue tension on PDMS rods was significantly higher in fibrotic tissues (red) when compared to control samples (black) (average ± SEM, two-way ANOVA p = 0.002 between treatments, N = 5). No significant changes were observed over time within each treatment. **d**, Maximum force of contraction (measured at peak cardiomyocyte contraction) was significantly lower in fibrotic tissues (average ± SEM, two-way ANOVA p = 0.0009 for D14, p = <0.0001 for day 21, N = 5). Force increased over time within control tissues (two-way ANOVA p = 0.022). **e**, Excitation threshold at day 21 was significantly higher in fibrotic tissues compared to controls, indicating poorer electrical connectivity (average ± SEM, p = 0.003, N = 7). **f**, Analysis of BNP secretion by ELISA shows fibrotic tissues have significantly higher BNP secretion than controls (average ± SEM, p < 0.0001, N = 4).

Analysis of mRNA in the samples at endpoint revealed significant changes in RNA expression, with approximately 1,000 genes showing differential expression (**Supplemental Fig. 5a**, p ≤ 0.05, ≥2-fold). In line with a fibrotic tissue profile (Frangogiannis, 2019; Travers et al., 2016), there was an upregulation in the extracellular matrix proteins collagen type I (*COL1A2*), fibronectin (*FN1*) and periostin (*POSTN*), as well as induction of vimentin (*VIM)*, TGFβ1 (*TGFB1*), α-SMA (*ACTA2*), lysyl oxidase (*LOX*), and connective tissue growth factor (C*CN2*) (**Fig. 3a**). Gene set enrichment analysis showed corresponding gene set upregulation, including extracellular matrix, extracellular structure, collagen binding, and collagen fibril organization (**Fig. 3b**).

**Figure 3.**
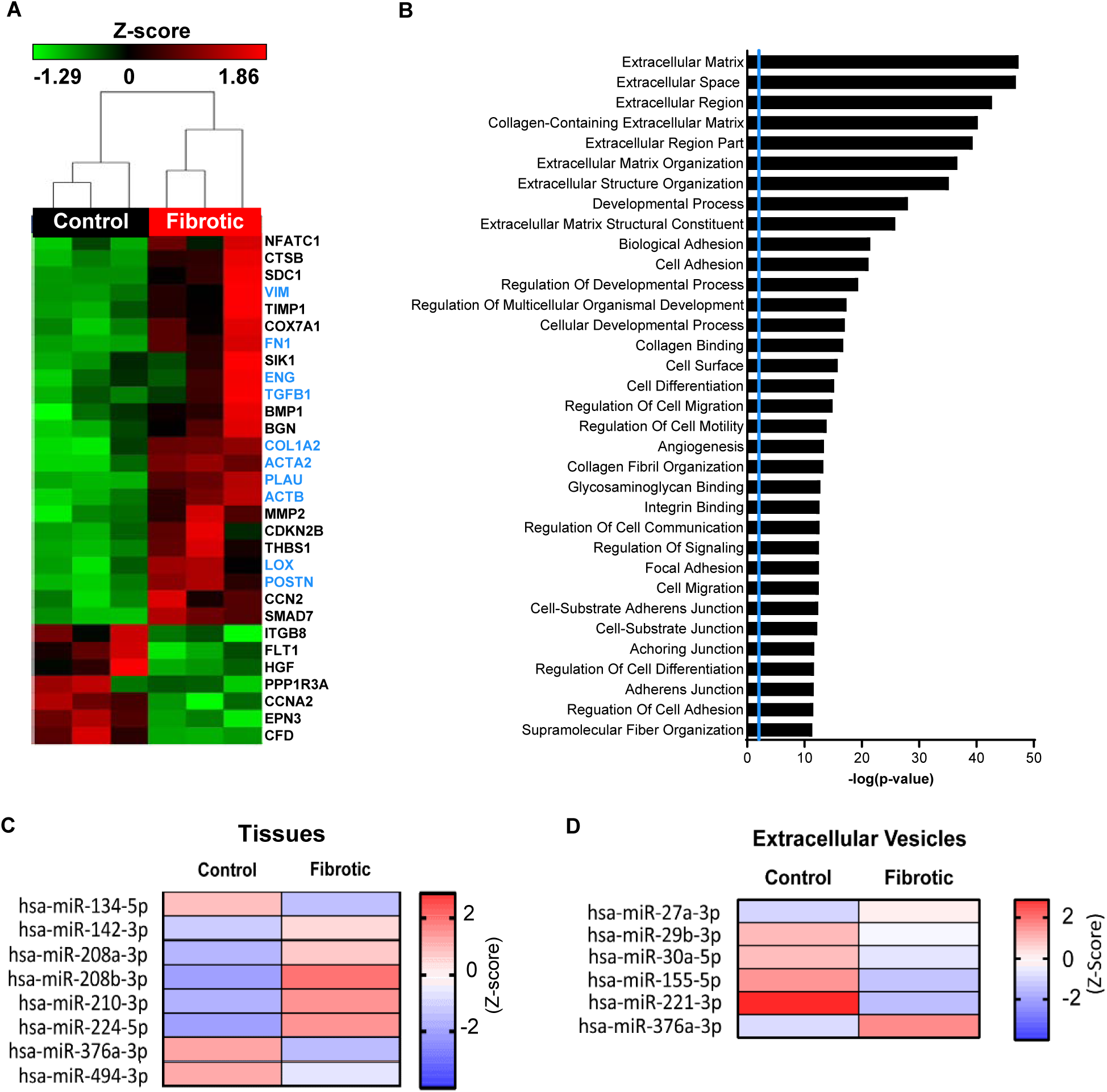
Comparative transcriptomic analysis between control and fibrotic tissues shows recapitulation of a fibrotic signature. **a**, Heat map showing unsupervised clustering of selected fibrosis-relevant genes that were differentially expressed (p ≤ 0.05, ≥2-fold) in the fibrotic tissues compared to the control tissues, z-scores for the RPKM. **b**, Gene set enrichment analysis (GSEAs) performed in 3 independent experiments showing enrichment for genes involved in cardiac fibrosis (-log transformation of significant genes at p≤0.05, threshold line (blue) at p = 0.05). **c**, Heat map depicting up-regulation or down-regulation of the miRs associated with cardiac fibrosis in the fibrotic tissues compared to the control tissues. **d**, Heat map showing changes in extracellular vesicles-associated miRs of fibrotic tissues compared to controls (p<0.05 and fold change of ≥2-fold). N = 3 for all.

Given their potential as biomarkers and as treatment targets, we also investigated the expression of cell-intrinsic and secreted microRNAs (miRs) in our hCF-on-a-chip. Interestingly, several miRs were significantly differentially expressed in fibrotic tissues compared to controls, including miRs -142 and -208 (**Fig. 3c** and **Supplemental Table 1**), which have been previously found to be dysregulated in the setting of cardiac dysfunction and tissue fibrosis (Su et al., 2015; van Rooij et al., 2007; Voellenkle et al., 2010; Zhu et al., 2013). KEGG pathway analysis of signaling pathways that are predicted to be targeted by differentially expressed miRs revealed pathways related to cardiac fibrosis and heart failure, such as: extracellular matrix-receptor interactions (Bonnans et al., 2014), PI3K-Akt (Travers et al., 2016), TGF-β signaling (Khalil et al., 2017; Meng et al., 2016), Hippo signaling (Leach et al., 2017), mTOR (Gonzalez-Teran et al., 2016), Gap junctions (Severs et al., 2004), and focal adhesion (Dalla Costa et al., 2010) pathways (**Supplemental Fig. 6a**). Moreover, although no differences in size were observed in extracellular vesicles secreted into the media size (**Supplemental Fig. 6b**), analysis of miR content in extracellular vesicles revealed significant differences between control and fibrotic samples, including increases in miR-376a and decreases in microRs-27a, -29b, -30a, -155 and - 221 (**Fig. 3d** and **Supplemental Table 2**).

### Testing the suitability of the model to study drug efficacy

Since BNP secretion is the best prognostic indicator in heart failure, we tested the effect of standard-of-care drugs carvedilol and losartan on BNP secretion as proof of concept for the utility of the platform in drug studies. Given that a clinical diagnosis is usually reached at a time when cardiac functional outcome is impaired, we elected to treat the fibrotic tissues at a time point when tissues were fully compacted and force of contraction was already significantly decreased compared to controls. Therefore, drug treatment started 2 weeks post-seeding (day 14), when fibrotic tissues were randomized into treatment groups (**Fig. 4a**). In line with their use in the clinic due to their efficacy, fibrotic tissues treated with either drug significantly lowered BNP secretion (**Supplemental Fig. 7a, b**). Importantly, neither drugs showed significant effects on BNP secretion if tested in 2D monolayers at the same dose (**Supplemental Fig. 7c**, p=0.09 and p=0.17 for losartan and carvediolol, respectively), suggesting that recapitulating the higher order intercellular interaction is likely important to study the biological processes that take place *in vivo* and in response to drugs. Moreover, neither drugs affected either passive tension or active force (**Supplemental Fig. 8**) irrespective of if the treatment was performed for 1 or 2 weeks.

**Figure 4.**
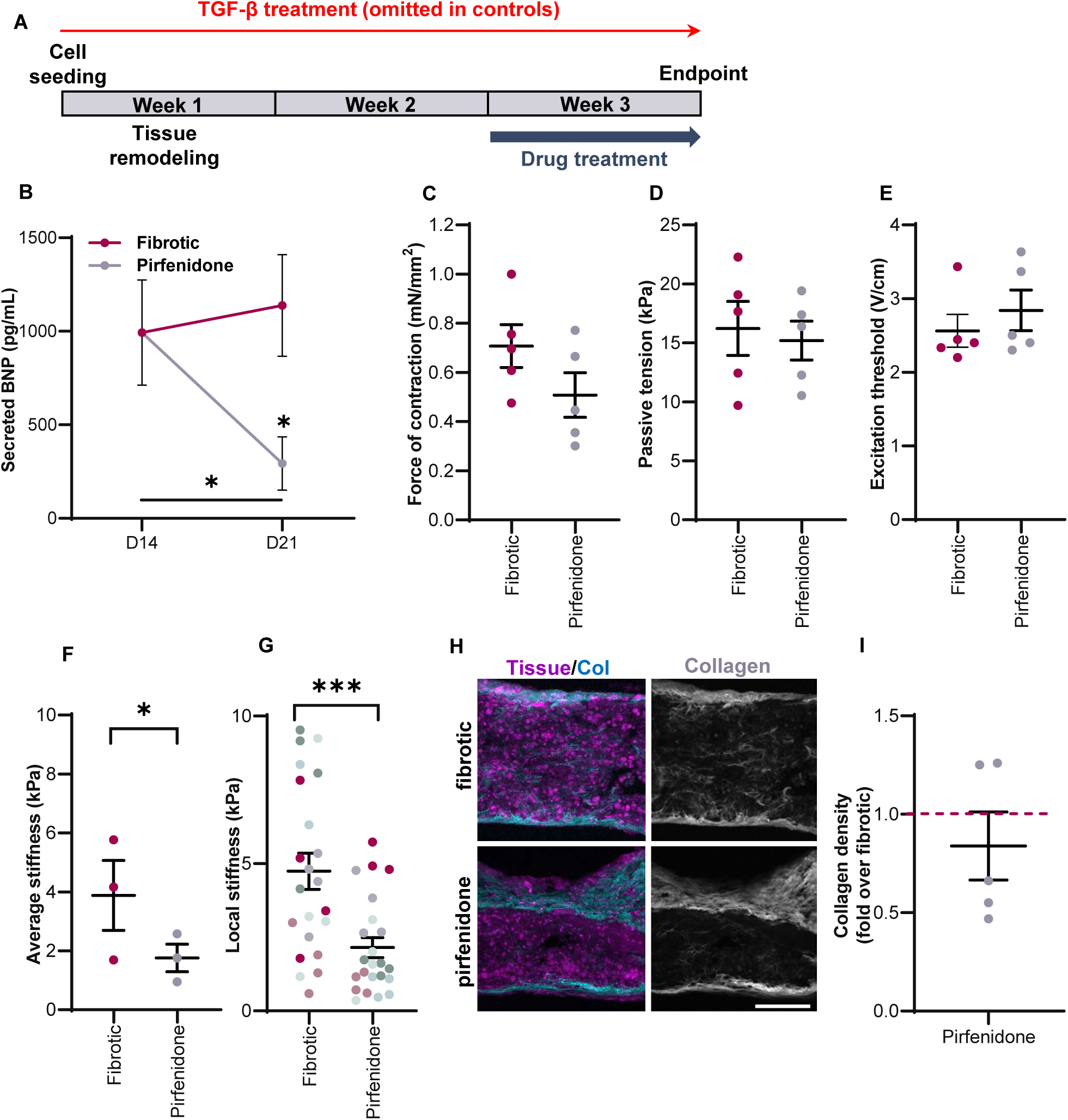
Treatment with the anti-fibrotic drug pirfenidone significantly decreases tissue stiffness and BNP secretion but does not alter other parameters. **a**, Schematic of the drug treatment timeline. Fibrotic samples were randomized into treatment or not with pirfenidone (2.5 mM) at the end of week 2 in culture. **b**, Treatment with pirfenidone (grey) significantly decreases BNP secretion compared to untreated fibrotic tissues (red) (average ± SEM, two-way ANOVA p= 0.031 D21 pirfenidone vs fibrotic, p = 0.048 D14 vs D21 pirfenidone, N = 4). **c**, No differences were observed in the force of contraction (measured at peak contraction) nor (**d**) passive tension between fibrotic and pirfenidone-treated tissues at day 21 (average ± SEM, N = 5). **e**, Excitation threshold between fibrotic and pirfenidone-treated tissues at day 21 (average ± SEM, N = 5). **f**, Average tissue stiffness assessed by atomic force microscopy showed a significant decrease in pirfenidone-treated compared to fibrotic tissues at day 21 (average ± SEM, p= 0.013, N = 3). **g**, Individual stiffness for each point probed shows reduced heterogeneity in pirfenidone-treated tissues compared to fibrotic (N = 3, n = 6 tissues total, 4 points probed/tissue). Each dot represents different points probed in the same tissue represented by same color (average ± SEM, p = 0.0005). **h**, Representative images of fibrotic and pirfenidone-treated tissues assessed by second harmonic generation. Collagen (Col), cyan (left) and grey (right). Cells, magenta (auto-fluorescence). Scale bars, 250 μm. **i**, Collagen area over tissue area quantification of fibrillar collagen deposition in fibrotic of pirfenidone-treated samples at day 21 by second harmonic generation vs control (dashed red line) (average ± SEM, N = 5).

We also tested the potential application of pirfenidone, a drug used in the treatment of idiopathic pulmonary fibrosis (Nanthakumar et al., 2015; Noble et al., 2011) that also had positive effects in pre-clinical animal models of cardiac fibrosis (Mirkovic et al., 2002; Wang et al., 2013). We chose the lowest pirfenidone dose tested that induced the highest decrease in BNP secretion (**Supplemental Fig. 9a**) with no cytotoxicity based on liver studies (Westra et al., 2014). Tissues were treated for 1 week and assessed at endpoint (day 21, **Fig. 4a**) for collagen deposition, tissue stiffness, passive and active force, and BNP secretion. The miR signature in the tissues and in secreted extracellular vesicles was also assessed.

Tissues treated with pirfenidone showed a dose-dependent decrease in secreted BNP (**Supplemental Fig. 9a**). This was due to a decrease in BNP secretion relative to fibrotic tissues, as BNP levels were already high at the time of treatment (**Fig. 4b**, day 14). Of note, while BNP secretion following pirfenidone treatments of TGF-β1-stimulated co-cultures in 2-dimensional (2D) culture was 26% lower relative to the fibrotic group, it was reduced by 74% relative to the fibrotic group in the 3-dimensional (3D) tissues (**Supplemental Fig. 9b**), suggesting modulation of 3D-specific effects on cardiomyocyte stress. Although no significant changes were observed in force of contraction, passive tension, or excitation threshold after treatment with pirfenidone for one week (**Fig. 4c, d, e** and **Supplemental Video 5**), there was a significant decrease in average and local tissue stiffness (**Fig. 4f, g**). However, quantification of collagen deposition by second harmonic generation imaging revealed that collagen content was not significantly affected by treatment with pirfenidone (**Fig. 4h, i**). While we used one hiPSC-CM cell line for these assessments, an additional run with an independent cell line showed the observed trend for BNP secretion, force of contraction, passive tension, and excitation threshold for control, fibrotic, and pirfenidone-treated tissues at day 21 (**Supplemental Fig. 10**).

At day 21, approximately 500 genes showed differential expression between fibrotic and pirfenidone-treated tissues (**Supplemental Fig. 11**, p < 0.05, ≥2-fold). Of particular interest, pirfenidone-treated tissues showed significant downregulation of cardiac fibrosis markers including periostin, lysyl oxidase, connective tissue growth factor, and collagen type I (**Fig. 5a**). Although imaging did not reveal decreased collagen content, the decreased collagen type I mRNA expression at day 21 suggests a tendency for decreased collagen deposition (**Fig. 5a**). Moreover, the decreased lysyl oxidase expression (**Fig. 5a**) suggests there is a reduction in collagen crosslinking, which is consistent with the decrease in tissue stiffness after pirfenidone treatment (**Fig. 4f, g**). Curiously, data also revealed an upregulation of α-SMA and fibronectin.

**Figure 5.**
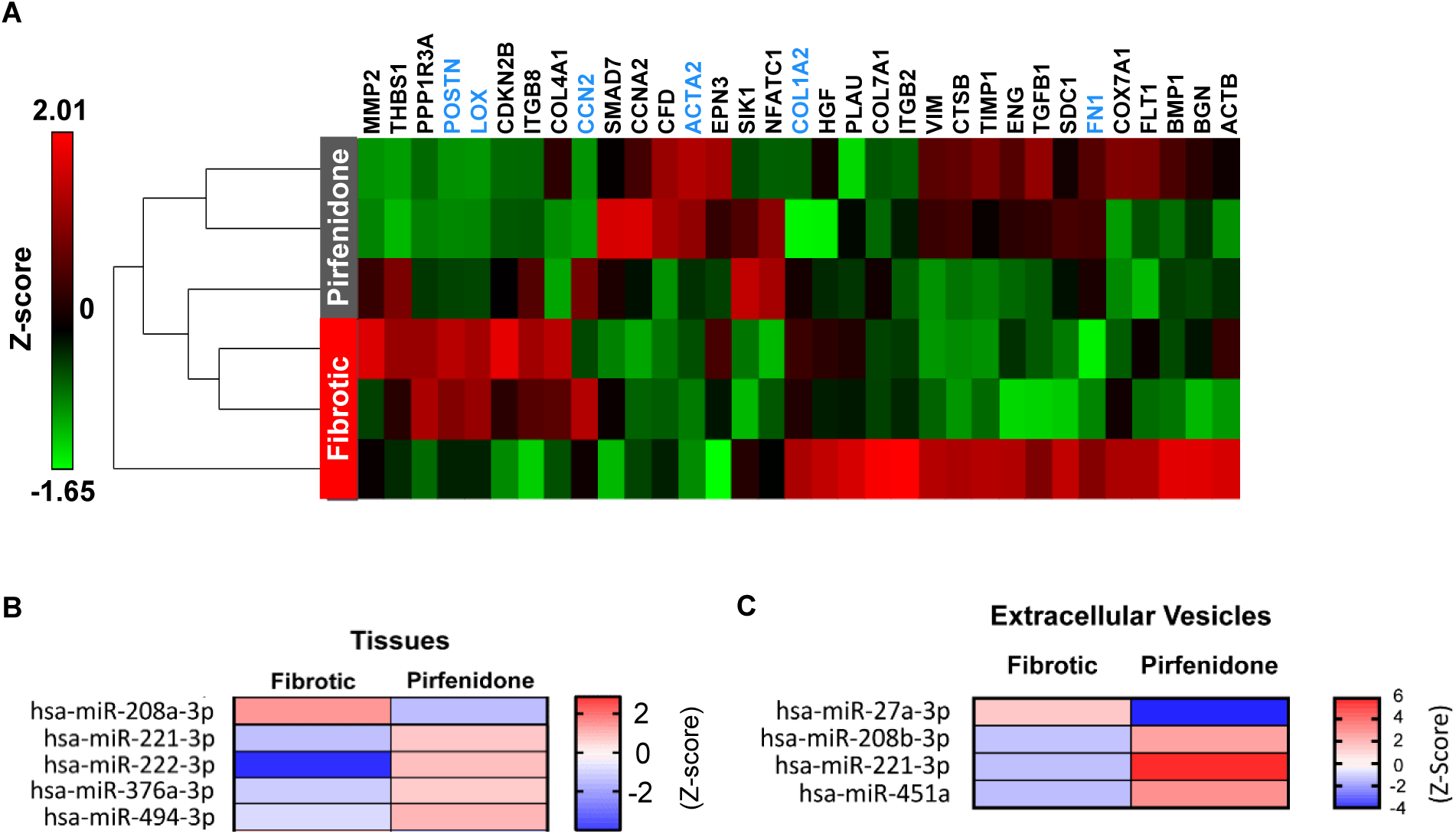
Comparative transcriptomic analysis between fibrotic and pirfenidone-treated tissues show significant differences in accordance with improvement in fibrosis. **a**, Heat map showing unsupervised clustering of selected fibrosis-relevant genes that were significantly different (p ≤ 0.05, ≥2-fold) in the fibrotic tissues compared to the pirfenidone-treated tissues, z-scores for the RPKM. **b**, Heat map depicting differentially expressed miRs associated with cardiac fibrosis in the fibrotic tissues compared to the pirfenidone-treated tissues. **c**, Heat map showing changes in extracellular vesicles-associated miRs of fibrotic tissues compared to pirfenidone-treated (p<0.05, ≥2-fold). N = 3 for all.

Encouraged by the positive effects of pirfenidone on BNP secretion, tissue stiffness and transcriptome, we investigated if performing longer term drug treatment by starting at an earlier time point would have an impact in function. Interestingly, we observed that treating tissues for 2 weeks significantly decreased tissue passive tension but had no effect on force of contraction (**Supplemental Fig. 12**).

Analysis of miR expression in human cardiac tissues revealed that fibrotic tissues responded to pirfenidone treatment by significantly changing the expression of several microRNAs (**Supplemental Table 3**). Changes included the downregulation of miR-208 and upregulation of miRs -221, -222, -376a and -494 (**Fig. 5b**). KEGG pathway analysis of signaling pathways predicted to be affected by miRs that were significantly up- or down-regulated post-treatment with pirfenidone included pathways relevant to cardiac fibrosis and heart failure (**Supplementary Fig. 13a**). Interestingly, analysis of miR content in extracellular vesicles secreted into the media revealed significant differences with drug treatment, including increase in miRs-208b, -221 and 451a, and a decrease in miR-27a (**Fig. 5c** and **Supplemental Table 4**). No differences in extracellular vesicles size were observed (**Supplemental Fig. 13b**).

## Discussion

We describe for the first time the development of a human heart failure model that is driven by cardiac fibrosis and displays the major hallmarks of the disease, including increased collagen deposition, higher tissue stiffness, loss of contractile function, and induced BNP secretion. Moreover, we demonstrated that the mRNA and miR signatures of fibrotic tissues recapitulate those of samples from human patients with cardiac fibrosis and/or heart failure (Thum, 2014).

Previous methods for studying cardiac fibrosis relied on 2D models (monolayers) in plastic substrates that fail to recapitulate the biomechanical properties and tissue-level organization (alignment, three-dimensionality, extracellular matrix, cell heterogeneity) of cardiac tissue, or in 3D systems without tissue alignment or key functional assessment capabilities such as force of contraction (Lee et al., 2019; Sadeghi et al., 2017). Here, we performed a comprehensive assessment of multiple key aspects of cardiac fibrosis using human cells with the concomitant assessment of function in real time, and the screening of relevant drugs. This simplified model more closely resembles the native tissue architecture and could allow for added complexity by including additional cells types (i.e. endothelial cells, immune cells) that may be relevant for disease progression. This will help to dissect the role of different cell types and signaling pathways in disease pathophysiology.

In the drug testing front, treatment of fibrotic tissues with standard-of-care drugs decreased BNP secretion but did not have any effects in either passive or active force at the time points treated. Meanwhile, despite not being able to fully reverse cardiac fibrosis (evidenced by lack of change in collagen deposition and failure to restore function), our findings showed significant beneficial effects of treatment with pirfenidone including decreased tissue stiffness, downregulation of key fibrosis-related genes, reversal of a fibrotic miR signature, decrease in passive force, and reduced BNP secretion. Secreted BNP levels are the gold-standard diagnostic marker for detecting cardiac events and have been found in multiple clinical studies to be the best prognostic indicator in heart failure (Troughton et al., 2014), with patients presenting low plasma BNP concentrations having the best outcomes (Aspromonte et al., 2008; Bettencourt et al., 2004; Wong et al., 2004). By this metric, our findings indicate that our hCF-on-a-chip model is superior to 2D cultures, and represents a novel functional approach for drug screening with improved physiological relevance, accuracy and throughput. It also indicates that treatment with pirfenidone may be beneficial for patients with cardiac fibrosis and likely not toxic for human cardiomyocytes, although a larger number of hiPSC lines would be needed to further support this observation. Despite many positive effects, pirfenidone did not reverse loss of contractile force, suggesting that it aids to prevent further decline in the presence of the fibrotic stimulus (TGF-β1), but is not able to fully reverse pathological changes in force generation at the dose and time tested. This is in agreement with the cardioprotection conferred by pirfenidone treatment in animal models of cardiac fibrosis (Wang et al., 2013; Yamagami et al., 2015). Moreover, despite not affecting contractile force, longer treatments with pirfenidone led to an improvement in passive force which can positively impact cardiac function.

While the approval of pirfenidone for clinical application is a major advance in fibrosis therapy, its use is restricted to the treatment of IPF patients and its mechanism of action is not fully understood. By creating a disease-relevant model, the hCF-on-a-chip technology may enable target validation and drug repositioning by providing a rapid assessment of functional outcomes in a human-relevant model. Moreover, present findings suggest this platform can help to inform drug application and safety, as well as tissue-specific roles (i.e. cardiac-specific toxicity) which could accelerate drug repositioning and save billions of dollars. It also allows for the study of the progression of fibrotic remodeling in a more complex (multi-cell type), 3D environment and can be used with patient-specific iPSC-derived cells, enabling personalized disease modeling and drug testing.

Although there has been an increase in the search for biomarkers to predict disease detection and prognosis or to allow disease stage stratification, a comprehensive understanding of the role of these molecules in disease pathophysiology is lacking (van Rooij and Olson, 2009). In the case of miRs, it is difficult to predict from human studies if a given miR is truly causative of the disease, or if it is instead modulated as a response to functional cardiac impairment, or if it has a role in attenuating disease progression (i.e. an attempt to rescue function). The technology described here will aid in elucidating some of these aspects. For instance, despite being significantly increased in fibrotic tissues, the expression of miR-208, a cardiac-specific miR that was shown to be necessary for cardiomyocyte hypertrophy and cardiac fibrosis (van Rooij et al., 2007), did not significantly change in secreted extracellular vesicles in fibrotic samples. This suggests that cell-intrinsic tissue and secreted miRs may have different roles and that secreted miRs may have specific function, as proposed elsewhere (Tkach and Thery, 2016). It also suggests that the loading of particular miRs into vesicles is a highly specific and regulated process. In support of this, we saw that miR-221, found to target TGF-β-mediated pro-fibrotic SMAD2 (mothers against decapentaplegic homolog 2) signaling and its downstream gene expression (Verjans et al., 2018), was not differentially expressed in control or fibrotic tissues but was significantly downregulated in secreted extracellular vesicles in fibrotic samples. Interestingly, treatment with the anti-fibrotic drug pirfenidone led to a 2.1-fold increase in the miR-221 expression in the hCF-chip tissues (**Supplemental Table 1**), and an even higher increase in extracellular vesicles (32-fold). This is in line with its recently described role in counteracting myocardial fibrosis in pressure overload-induced heart failure by blunting TGF-β-induced pro-fibrotic signaling (Verjans et al., 2018) and suggests a possible new mechanism of action for the drug. It also demonstrates the potential of the human hCF-chip platform to validate animal findings in a human-relevant model.

A growing body of evidence demonstrates extracellular vesicles are a universal cell feature and potential candidates for cellular gene transfer (Sluijter et al., 2014). However, the biology and biogenesis of shed extracellular vesicles are not completely understood (Tricarico et al., 2017). This includes a lack of a basic understanding of how particular miRs are preferentially loaded into vesicles, and which direction this vesicle-mediated cell communication occurs (signal-sending vs. -receiving cell types), and how an architectural complex environment composed of multiple cell types and with relevant biomechanical cues may play a role in vesicle signaling. Therefore, there is potential for application of the hCF-on-a-chip platform in the study of miR function which should increase our understanding of these messengers and improve adoption of biomarkers and development of miR-specific target drugs.

In conclusion, we have developed a human cardiac fibrosis-on-a-chip (hCF-on-a-chip) platform that recapitulated the distinctive features of cardiac fibrosis and provided proof-of-principle for using this platform for multi-parameter, phenotypic analysis of a drug recently approved by the FDA for IPF treatment. We utilized the human hCF-on-a-chip model to screen for possible biomarkers of human cardiac fibrosis, including secreted extracellular vesicles and replicated some trends already described in the literature, including in human clinical samples. We expect that the ability to increase the complexity of these tissues and the pathophysiologically relevant capabilities of the human hCF-on-a-chip devices, including cardiomyocyte functional analysis, will expedite the understanding of disease progression and will uncover alternative targets to treat heart failure and cardiac fibrosis.

## Experimental Procedures

### Device manufacture

Devices were designed using Autodesk Fusion 360 and milled from poly(methylmethacrylate) (PMMA) or polystyrene using a CNC milling machine (Personal CNC 770, Tormach, WI, USA). Polydimethylsiloxane (PDMS) (Dow Corning Corporation, cat# 3097358-1004) was mixed at indicated ratios of curing agent to PDMS base to produce the prepolymer mixture, degassed in a vacuum chamber, and then used to fill 27-gauge syringe needles. The prepolymer mixture was cured at 80°C for 2 hours and then extracted from the needles, yielding PDMS rods 210 μm in diameter. The resulting rods were placed in pairs horizontally in supporting wells in the PMMA or polystyrene devices and further secured outside the wells with approximately 0.2 uL 1:10 PDMS cured overnight.

### Cell culture and seeding

Human ventricular cardiac fibroblasts (Lonza cat# CC-2904) were grown in 1% gelatin coated flasks in DMEM (ThermoFisher cat#11885084) supplemented with 20% fetal bovine serum (FBS). Fibroblasts up to passage 6 were used. hiPSC-CMs (iCell lines A and B – **Supplemental Table 5**, cat# CMC 100 010 001 and R1106, >95% purity) were thawed and immediately used for seeding. Cells were seeded in fibrin gels containing a mixture of fibrinogen (5mg/mL), aprotinin (0.00825 mg/mL), thrombin (2 U/mL; Sigma-Aldrich cat# T7513), Matrigel (1 mg/mL, (VWR, cat# 354230), Y-27632 (10 nm, STEMCELL cat# 72304), in DMEM. A total of 250,000 cells were seeded in 7 uL of gel/device well. Human fibroblasts and hiPSC-CM were seeded at a 1:3 ratio. Gels were allowed to polymerize for 45 minutes to 1 hour at 37°C. For fibrotic conditions, cardiac fibroblasts were pre-treated in plastic tissue-culture flasks with 10 nM human TGF-β1 (Cell Signaling Technology, cat# 8915LC) for 48h. Tissues were cultured in iCell plating media (iCell CMM-100-010-001) with 1X penicillin/streptomycin/Amphotericin B (P/S/A, Gibco cat# 15240062). After 48h, the plating media was replaced with StemPro-34 Serum Free Media (ThermoFisher cat# 10639011) supplemented with StemPro-34 Nutrient Supplement, L-glutamine (2mM), L-ascorbic acid (50ug/mL), transferrin, 1-thioglycerol (0.039ul/mL, Sigma-Aldrich cat# M6145), and 1X P/S/A. Media was replaced every second or third day for the full duration of the experiment. The media for fibrotic tissues was also supplemented with 10 nM TGF-β1 (or vehicle for controls) for every media replacement.

Treatment with drugs, including pirfenidone (Milipore-Sigma, cat# P2116), losartan (Milipore-Sigma, cat# 61188), and carvedilol (Milipore-Sigma, cat# C3993) was performed during the second and third, or only third week of culture concomitantly to 10 nM TGF-β1.

For monolayer culture BNP assessment, cardiac fibroblasts and hiPSC-CMs were combined and plated on gelatin-coated plates at a 1:3 ratio. Media type was the same as with the tissues, and fibrotic sets were similarly pre-treated and maintained in 10 nM TGF-β1. Drug treatments with pirfenidone, losartan, and carvedilol were done two days after cell seeding, and media was collected on day four.

### Electrical stimulation

For electrical stimulation, devices were placed between two parallel carbon rods (Ladd Research, cat# 30250) spaced 1.5 cm apart inside a petri dish, each with a diameter of 3 mm. Platinum wire, a biocompatible material, was securely wrapped around each carbon rod and extended to the outside of the petri dish. Each platinum wire was then connected to a pulsed electric stimulator (Astro-Med Grass S88X Stimulator). Excitation threshold was assessed after a fresh media change. Petri dishes were placed under a microscope, and voltage was gradually increased from 1 V until the tissue changed its beating pattern to match the 1 Hz pulses.

### Force assessment in the tissues

Video recordings of the rod deflection were captured under electrical stimulation using a Leica EC3 camera. Image analysis was performed with ImageJ 1.51n. Tissue widths were measured at the middle of the tissue (reported in **Fig. 1b**) and at the PDMS rods (used for force calculations). Rod deflection for passive force was measured as the distance between the PDMS rod in the tissue’s relaxed state, and the PDMS rod at non-deflected position. For force of contraction (active force), peak rod deflection under electrical stimulation at 1 Hz was measured. For active force calculations, passive force was subtracted from force calculated with the maximum deflection. Tissue widths and rod deflection measurements were performed by an experimentalist blinded to the conditions.

### Force calculations

Suspended weights were used to establish force-deflection curves, accounting for different tissue widths at the rods. Regression analysis was performed on the measurements using the MATLAB R2018a Curve Fitting Toolbox, from which we obtained the following equation:

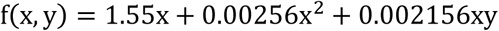

The function *f* represents force (N), while *x* is the PDMS rod deflection from its non-deflected position origin (m), and *y* is the tissue width at the midspan of the PDMS rod (m). As the model is non-linear, passive deflection force (or passive tension) on the PDMS rod as the tissue is relaxed must be subtracted from total force at maximum PDMS rod deflection during a tissue pulling in for a beat to find the active force of contraction (**Supplemental Fig. 2**).

For normalization by cross-sectional area for active force and passive tension, cross-sectional area for tissues was calculated as the area of an oval (π\**r*_1_* *r*_2_), where *r*_1_ is half of the tissues’ thickness, and *r*_2_ is half of each individual tissues’ width at its mid-span. Tissues’ mid-span width was measured under the same conditions as force. Tissue heights were measured in 4% paraformaldehyde (PFA)-fixed tissues on their sides, and an averaged value (300 μm) was used across all tissues.

### Assessment of tissue stiffness by atomic force microscopy

Live tissues were assessed with atomic force microscopy at endpoint. Tissues were taken out of device wells by the PDMS rods and secured in place on top of charge adhesive glass slides by using cover slips coated in high vacuum grease to hold PDMS rods down. Blebbistatin (10 μM) in PBS was added to prevent the tissues from contracting (beating) and to keep them hydrated. Measurements were taken with a Bruker BioScope Catalyst atomic force microscope using spherical tips manufactured by adding 5 μm radius spherical polystyrene beads to Bruker MLCT-O10 cantilevers using epoxy glue. The cantilevers had a 0.03 N/m nominal spring constant, but precise spring constants were calibrated before each experiment using the thermal tune method as per manufacturer instructions. Four spots were probed near the center of each tissue using the contact mode in fluid setting with a 5 nN trigger force and the linear region of each force curve was analyzed using the Hertz model following manufacturer protocols.

### BNP quantification

Levels of brain natriuretic peptide (BNP) secreted in the media were quantified by Human BNP ELISA Kit (Abcam, cat# ab193694) as per the manufacturer’s instructions. Media was collected on days 14 and 21 of culture. Cellular debris was spun down at 1500rpm for 10 min at 4°C, and the collected supernatant was kept frozen at -80°C prior to running the ELISA. For monolayers, media was collected on day 4 of culture.

### Tissue characterization

For analysis of tissue structure and quantification by immunohistochemistry, 4% paraformaldehyde (PFA)-fixed tissues were sectioned into into 14 μm thick slices. They were then stained for cardiac troponin T (ThermoFisher cat# MS-295, 1:100), vimentin (Cell Signaling cat# D21H3, 1:100), α-SMA (Sigma Aldrich cat# A2547, 1:500), phalloidin (A12379, 1:50), Hoechst 33342 (Sigma-Aldrich cat# B2261, 5µg/mL), collagen type I (SouthernBiotech cat# 1310-01, 1:250), collagen type III (SouthernBiotech cat# 1330-01, 1:250), ki67 (Abcam cat# ab9260, and BD Pharmingen cat# 550609, 1:100), and TUNEL (DeadEnd™ Fluorometric TUNEL System cat# G3250). Tissues were imaged at 20X magnification as z-stacks with an Olympus Fluoview FV1000 confocal microscope with imaging parameters kept constant for intensity quantification.

For second harmonic generation, tissues were fixed with 4% PFA and cryosectioned into 14 μm thick slices. Imaging was performed with a Zeiss LSM 710 NLO microscope.

Second harmonic generation collagen content was compared across samples by dividing the area of the collagen by the area of the general tissue (autofluoresence) using ImageJ. For connexin 43, α-SMA, collagen type I, and collagen type III, integrated intensity of SUM z-projected stacks was normalized over total nuclei. For ki67 and TUNEL quantification, total counts of ki67 and TUNEL-positive nuclei were divided over total nuclei counts. For cTnT and vimentin-positive counts, cTnT and vimentin-overlapping nuclei were divided over total nuclei counts.

### TRIzol RNA extraction

Three hundred microliters of TRIzol reagent was added to each tissue sample (2 mg fresh tissue weight) on ice, followed by 15–30 seconds of homogenization. Samples were incubated at RT for 5 minutes to allow for lysis and disruption of cells. To each tube 60 μL of chloroform was added, followed by 30 seconds of shaking/inversion and a 5-minute incubation at RT. Samples were then centrifuged at 17,000 × g for 15 minutes at 4 °C to separate RNA into aqueous phase. 120 μL of aqueous phase were carefully transferred to another microcentrifuge tube, and the RNA was precipitated with 120 μL of isopropanol at -20°C for 18 hours. Samples were centrifuged at 17,000 × g for 10 minutes at 4 °C to pellet RNA. Supernatant was removed, and the pellet was washed with 300 uL of 75% EtOH, followed by centrifugation at 17,000 × g for 5 minutes. The supernatant was carefully removed, and samples allowed to dry for 5 minutes at RT with the caps open. The RNA pellet was then resuspended in 20 μL of Ultrapure water and stored at -80 °C for downstream applications.

### Enrichment of extracellular vesicles from cell media

Approximately 15 mL of conditioned media was used as input for extracellular vesicle isolation. Conditioned media was centrifuged at 300xg, 4°C for 10 minutes, followed by an additional centrifugation at 2,000xg, 4°C for 15 minutes, to remove cells and cell debris. The media was then filtered through a 0.8µm filter (EMD Millipore, MA), with minimal pressure applied. The filtrate was then concentrated ∼60 times using 15mL 10kDa weight cut-off Amicon Ultra Centrifugal Filters (EMD Millipore) at 3,901xg, 4°C for 30 minutes. The particle size distribution and concentration of the concentrate was determined by nanoparticle tracking analysis using a NanoSight NS300 system (Malvern Instruments Ltd.), and the protein quantified using a Micro BCA Protein Assay Kit (ThermoFisher Scientific).

### Microfluidics based miR profiling

Reverse transcription was carried out with the Universal cDNA synthesis kit II (Exiqon) using a 16 µl reaction mixture containing 4 µL 5x reaction buffer, 9 µL nuclease-free water, 2 µL enzyme mix, and 1 µL synthetic U6 and Cel-miR-39 spike-in (Exigon). 20 ng of total RNA (5 ng/µL) was added to the reaction mixture and incubated as follows; 42°C for 60 min, 95°C for 5 min and then immediately cooled to 4°C. After RT, the cDNA product was diluted 1:10 in nuclease-free water and stored at −20°C until needed for amplification. Pre-amplification of cDNA was then initiated by creating a pool of 96 LNA™ miR Assays for each assay. The pre-PCR amplification reaction was performed in a 5 µl reaction mixture containing 2.5 µl TaqMan PreAmp Master Mix 2X (Applied Biosystems), 1.25 µl of 96-pooled LNA™ assay mix (0.25X) and 1.25 µl of diluted cDNA. The pre-amplification PCR was performed according to the following cycling conditions: one cycle 95°C for 10 min, 15 cycles at 95°C for 15 sec and then 60°C for 4 min. After pre-amplification PCR, the product was then diluted 1:10 in nuclease-free water and stored at −80°C until needed for amplification. Quantitative PCR of the miR targets was carried out using the 96.96 dynamic array (Fluidigm Corporation) following Exiqon’s recommended protocol (**Supplemental Table 6**). Briefly, a pre-sample mixture was prepared for each sample containing 275 µL of 2X TaqMan^®^ Gene Expression Master Mix (Applied Biosystems), 27.5 µL of 20X DNA Binding Dye Sample Loading Reagent (Fluidigm^®^ Corporation), 27.5 µL of EvaGreen™ (Biotium), and 82.5 µL 1X TE Buffer. 3.75 µL pre-sample mix was then combined with 1.25 µL each of diluted pre-amplified cDNA. Five microliters of unique assay mixes were prepared by combining 1.25 µL of an 1X LNA™ miR assay, 2.5 µL of 2X Assay Loading Reagent (Fluidigm^®^ Corporation), and 1.25µL 1X TE Buffer. The 96.96 Dynamic Array™ was primed with control line fluid in the integrated fluid circuit controller and 5 µL of both sample and assay mixes were loaded into the appropriate inlets. The chip was then loaded as well as mixed in the IFC controller, and then placed in the BioMark255 Instrument for PCR at 95°C for 10 min, followed by 35 cycles at 95°C for 10 sec and 60°C for 1 min. The data was analyzed with Real-Time PCR Analysis Software in the Biomark instrument (Fluidigm Corporation, CA) using a quality threshold of 0.65, linear (derivative) baseline correction method, and the auto (global) Ct threshold method. Following calibration and quality control of data, the global Ct mean of all expressed targets was used for data normalization.

### Determination of significant circulating miRs and pathway analysis

MicroRNAs were sorted based on up- or down-fold change regulation respectively and independently analyzed with DNA Intelligent Analysis (DIANA)-mirPath v3 (Vlachos et al., 2015) to determine over-represented biological pathways by computing p-values describing Kyoto Encyclopedia of Genes and Genomes (KEGG) pathway enrichment of predicted mRNA targets.

### RNA-Sequencing and data analysis

RNA was extracted from control, fibrotic, and pirfenidone treated tissues using the RNeasy Micro Kit (Qiagen Cat No. 74004) followed by RNA quality check on Agilent BioAnalyzer (Cat No. G2939BA) using the Agilent RNA 6000 nano kit (Cat No. 5067-1511). RNA samples with RNA integrity number (RIN) of 9-10 were chosen to prepare the cDNA libraries. Since the aim was to look for change in differential gene expression between the conditions, CATS mRNA-seq kit (with polyA selection) (Diagenode, Cat No. C05010051) was used. Briefly, the mRNA selection protocol was run on the chosen RNA samples as explained in the CATS mRNA-seq kit. Subsequently, cDNA libraries were prepared using 300 pg of mRNA after the selection.

Quality and size of the cDNA libraries were assessed using the High Sensitivity D1000 kit (Agilent, Cat No. 5067-5585) and run on Illumina Next-Seq 500 sequencing machine. The cDNA libraries were read using the 75 cycle kit V2.5 for single-read at 75 bp reading length and 35M sequencing depth. The data was analyzed using the partek flow software.

### Statistical Analysis

Statistical analysis was done with SigmaPlot 14.0 (Systat Software, Inc). Student’s t-test was used to assess significance between 2 groups. Mann-Whitney test was used to assess atomic force microscopy data. For conditions where a variable was analyzed over time, Two Way ANOVA was performed. A *p* value below 0.05 was considered significant. For miRs, data were normalized to the global mean expression of all miRs. GenEx 6 software (MultiD) was used for data pre-processing. Cq values were transformed to relative quantities and expression values were converted to log2 scale. Statistical Analysis was performed using both GenEx 6 software and GraphPad Prism 7 (GraphPad Software, Inc.) and a 2-fold cutoff was applied. Unless otherwise indicated, capital N’s indicate number of separate cell-seeding runs, while lower case n’s indicate number of tissues.

## Acknowledgements

This work was supported by grants from the Canadian Institutes of Health Research (CIHR), Institute of Circulatory and Respiratory Health (137352 and PJT153160) to S.S.N. A Discovery grant from the Natural Sciences and Engineering Research Council (NESERC)(RGPIN 06621-2017) to S.S.N. partially supported the following trainees: O.M., K.W. and M.K. O.M. was partially supported by a NSERC CREATE Training program in organ-on-a-chip engineering and entrepreneurship (TOeP) and NSERC Canada Graduate Scholarships-Master’s Program (CGS M). J.E.F. holds a Tier II Canada Research Chair in Vascular Cell and Molecular Biology and was supported by the CIHR-funded Canadian Vascular Network. D.G. was supported by a CIHR scholarship. B-U.M. was supported by CIHR 137352 to S.S.N., NSERC Discovery 436117-2013 and Engage EGP 500666-16 grants to E.W.K.Y.

## Author contribution

O.M. was the main experimentalist, M.K. performed video data analysis, R.H. contributed to cell culture assays and RNAseq analysis, B-U.M. contributed to development of the chip design, K.W. contributed to ELISA, A.Y.L.L., and Y.S. contributed to AFM assessments, D.G., J.E.F. and R.H. contributed to RNA and miR isolation, sequencing, and analysis, X.S. contributed to histology and statistical analysis, E.W.K.Y. contributed to chip design and fabrication, rod deflection characterization, and to manuscript writing, S.S.N designed the experiments, coordinated the project and contributed to the writing of the manuscript.

---

**Supplemental Figure 1.**
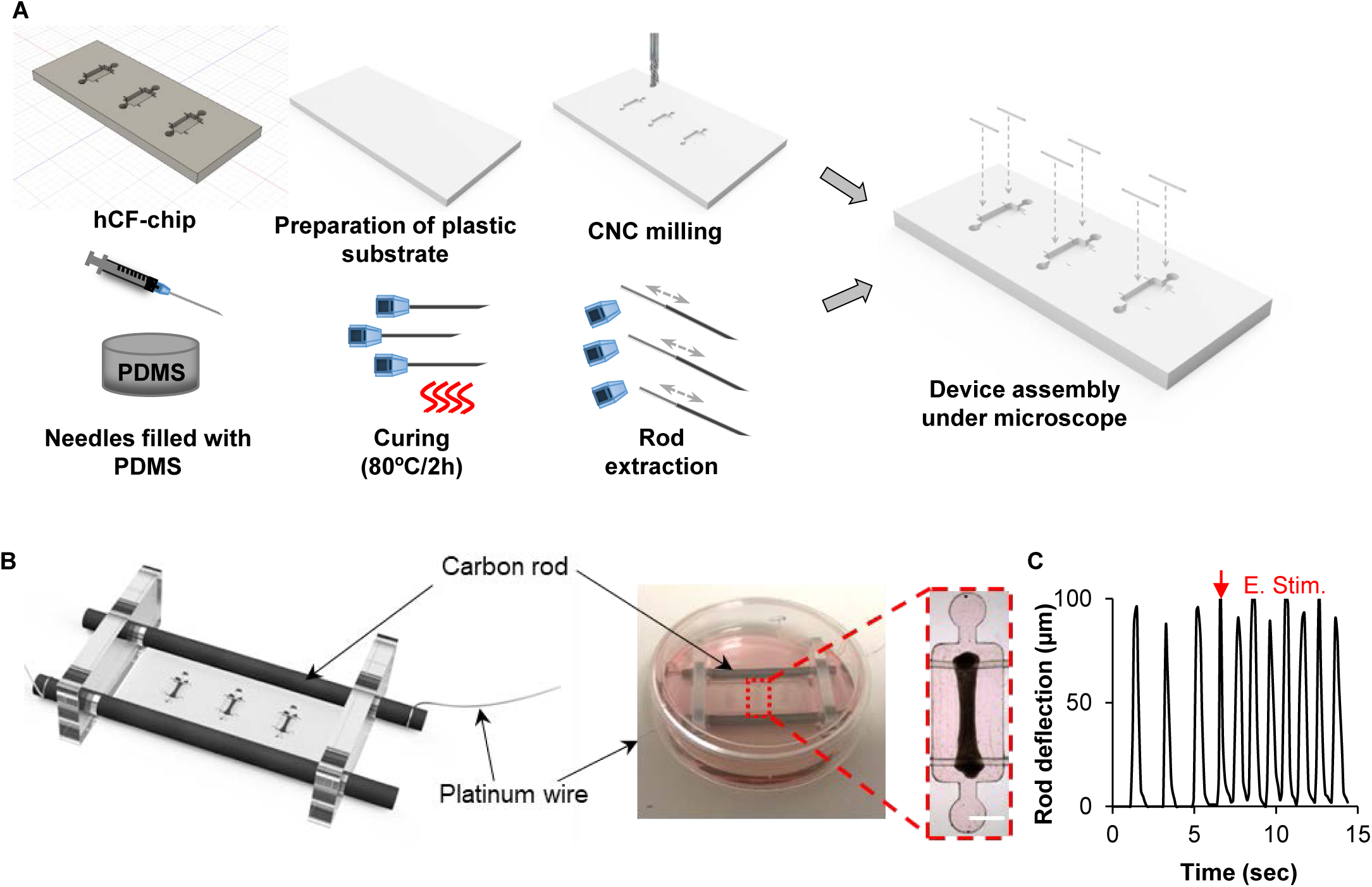
Platform design and fabrication. **a**, Microwells were generated on PMMA sheets by computer numerical control (CNC) milling (top panel). Elastomeric rods were obtained by using syringes to fill 27G gauge needles with PDMS prepolymer and curing agent. After curing, PDMS rods of ∼200 μm in diameter were pulled out of the needles and cut (bottom panel). Two-material chips were assembled by inserting the PDMS rods into the microwells. **b**, Schematic render of the device with microtissues (black) placed within an electrical field stimulation chamber (left). Picture of the device in the chamber (right). High magnification image of the heart-on-a-chip device (detail, top view, dashed line). Scale bar, 1 mm **c**, Rod deflection prior to and during electrical stimulation (E. stim.) at 1Hz frequency showed tissues can be paced.

**Supplemental Figure 2.**
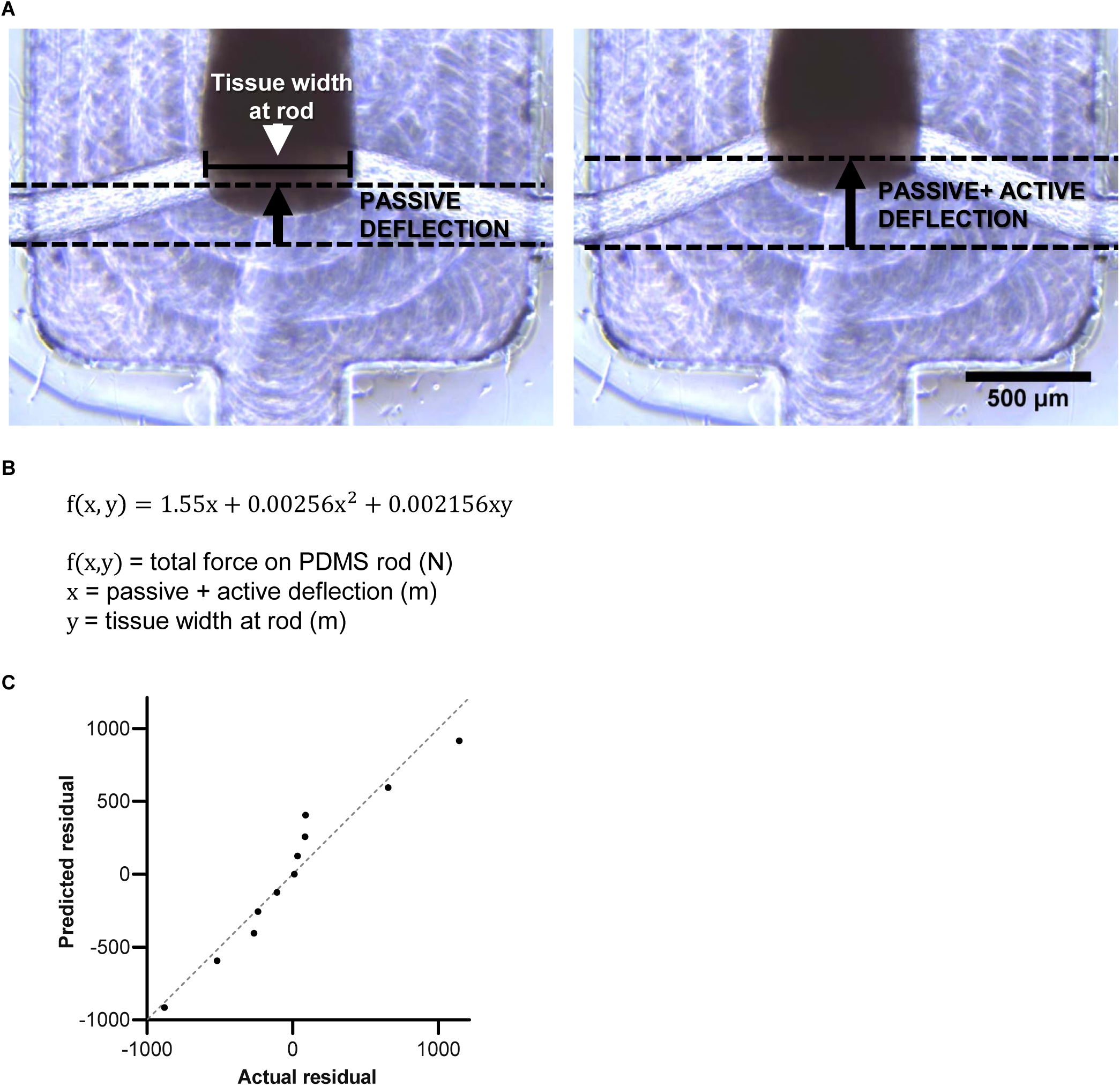
Force assessment in the chips. **a**, Representative images of rod deflection at relaxation (left) and peak cardiomyocyte contraction (right) in control sample at day 21 post-seeding. Tissue width at PDMS rods (white arrowhead) was measured as the edge-to-edge distance of the tissue construct that is pulling on the 1:30 PDMS rod. Rod deflection for passive tension (left image) was measured as the distance between the 1:30 PDMS rod in the tissue’s relaxed state (upper dashed line), and the rod at non-deflected state (bottom dashed line). For force of contraction (active force), peak rod deflection under electrical stimulation at 1Hz was measured (upper dashed line). For active force calculations, passive force was subtracted from force calculated with the maximum deflection. **b**, Equation used for determining total force exerted on the 1:30 PDMS rods by the tissues. **c**, Residuals plot for derived equation and validation set. Validation with three more sets of PDMS rods (N = 3, n = 4 per rod weights measured) resulted in an R^2^ value of 0.7645 for the equation.

**Supplemental Figure 3.**
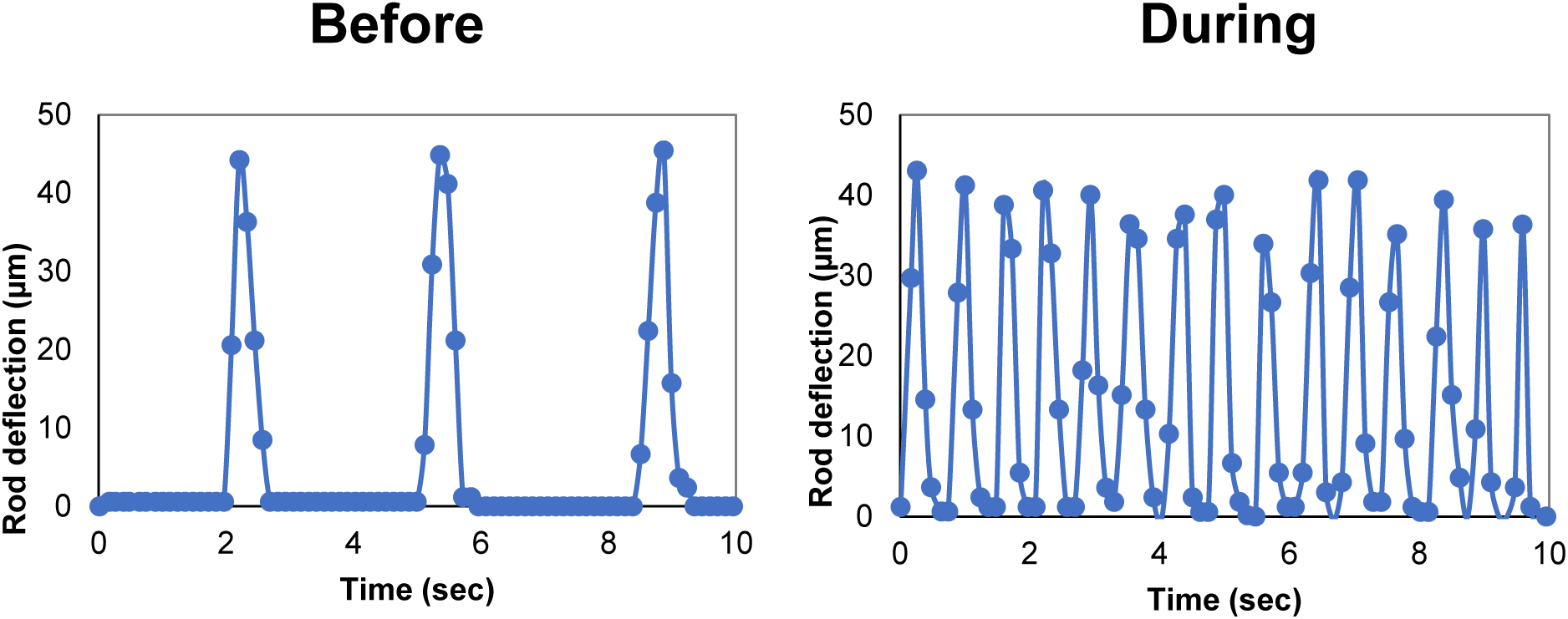
Tissues respond to isoproterenol treatment by increasing beating rate. Representative images of rod deflection at peak cardiomyocyte in spontaneous beating cardiomyoyctes (left) and in tissues treated with the beta-adrenergic receptor stimulator, isoproterenol (right).

**Supplemental Figure 4.**
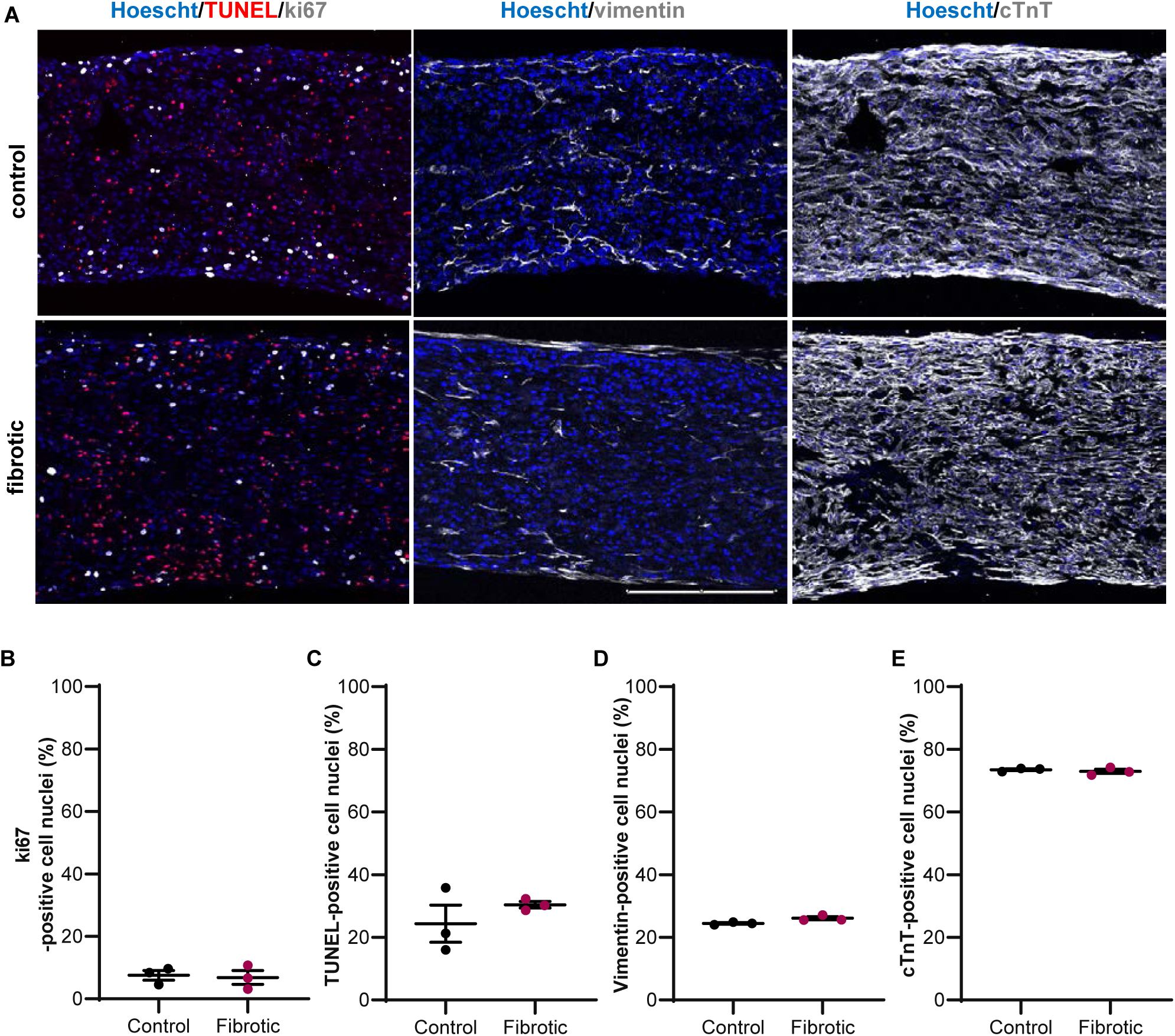
Quantification of viability at day 21 for control and fibrotic tissues. **a**, Representative immunostaining images for (left) TUNEL (red), ki67 (grey), (middle) vimentin (grey), and (right) cTnT (grey) for fibrotic and control tissues at day 21. Nuclear stain hoescht in blue. Scale bar, 250 μm. **b**, Cell proliferation assessment as percentage of ki67-positive nuclei. **c**, Terminal deoxynucleotidyl transferase dUTP nick end labeling (TUNEL) assay for percentage cell apoptosis. **d**, Percentage of nuclei overlapping with vimentin (fibroblast marker) staining over total nuclei. **e**, percentage of nuclei overlapping with cTnT staining (cardiomyocyte marker) over total nuclei. N = 3 for all, average ± SEM.

**Supplemental Figure 5.**
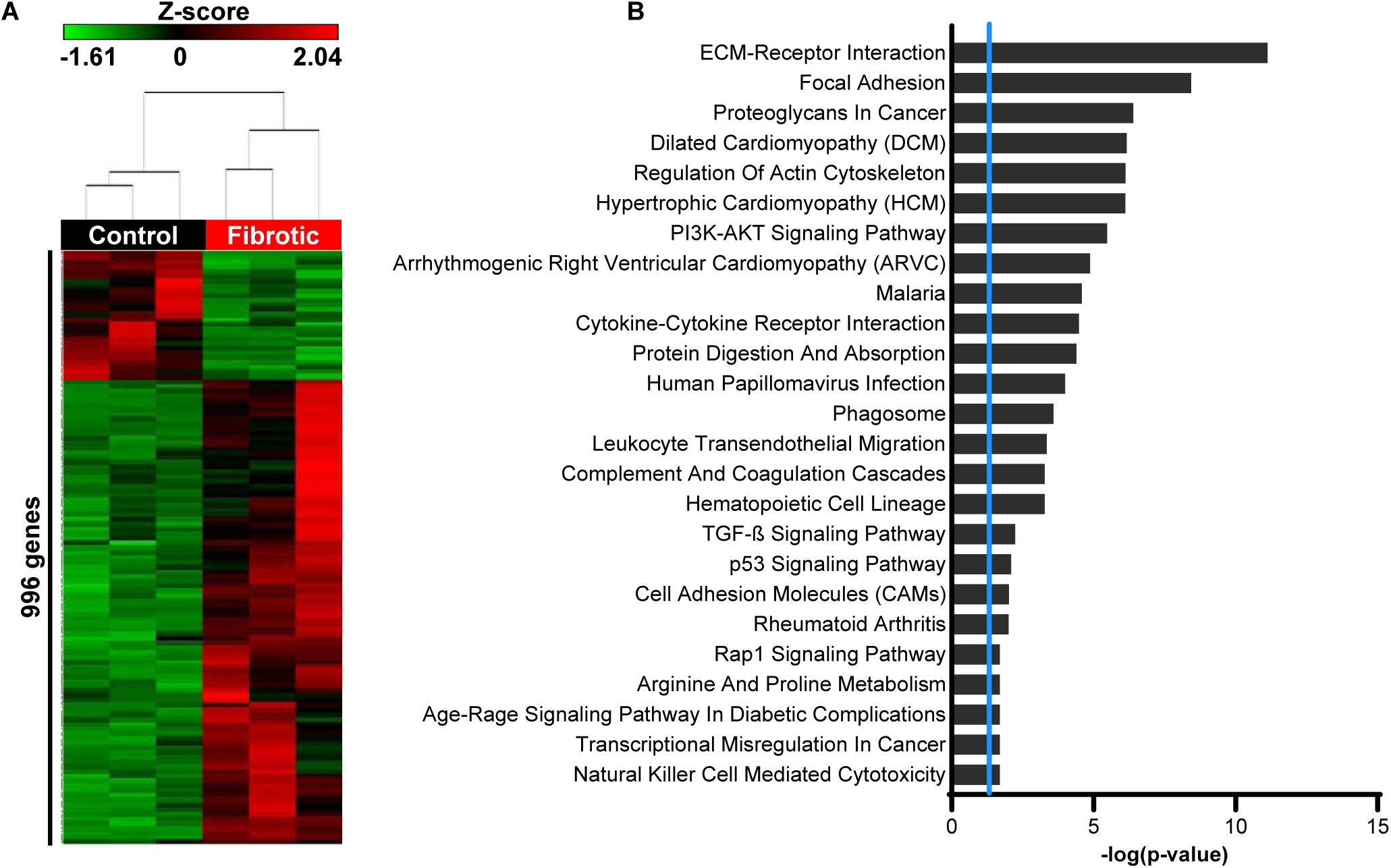
Transcriptomic analysis of control and fibrotic tissues by RNAseq. **a**, Unsupervised hierarchical cluster showing 996 genes differentially expressed between the fibrotic and control tissues, z-scores for the RPKM (p≤0.05, ≥2-fold, N = 3). **b**, Pathway analysis from sets of genes significantly upregulated in the fibrotic tissues compared to the control tissues based on the shared biological or functional properties as defined by a reference knowledge base, -log transformation of significant genes at p≤0.05 (threshold line (blue) at p = 0.01, N = 3).

**Supplemental Figure 6.**
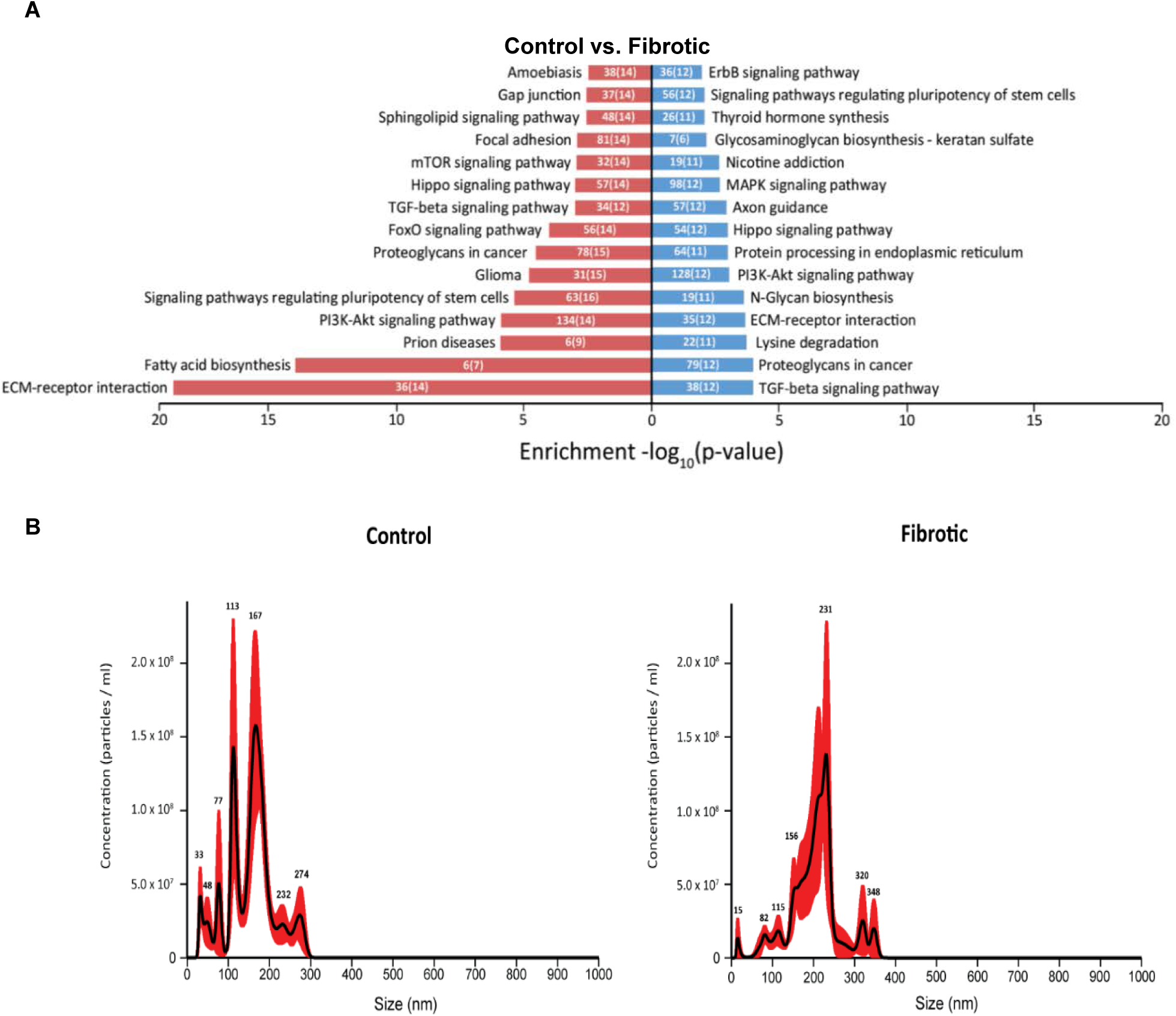
Human cardiac fibrosis-on-a-chip tissues exhibit a miR signature compatible with that of *in vivo* fibrosis. **a**, KEGG pathway analysis of signaling pathways predicted to be affected by miRs that were significantly down- (red) or up-(blue) regulated (N = 3). **b**, Representative size distribution of extracellular vesicles enriched from cell culture media for control and fibrotic tissues. The data were obtained from the binned averages of at least three 60 second NTA videos. The peak positions are marked. There are no statistical differences between groups for concentrations and 50 nm size bins. The black line represents the fitting curve of the binned data and the red error bars indicate ± SEM (N = 3).

**Supplemental Figure 7.**
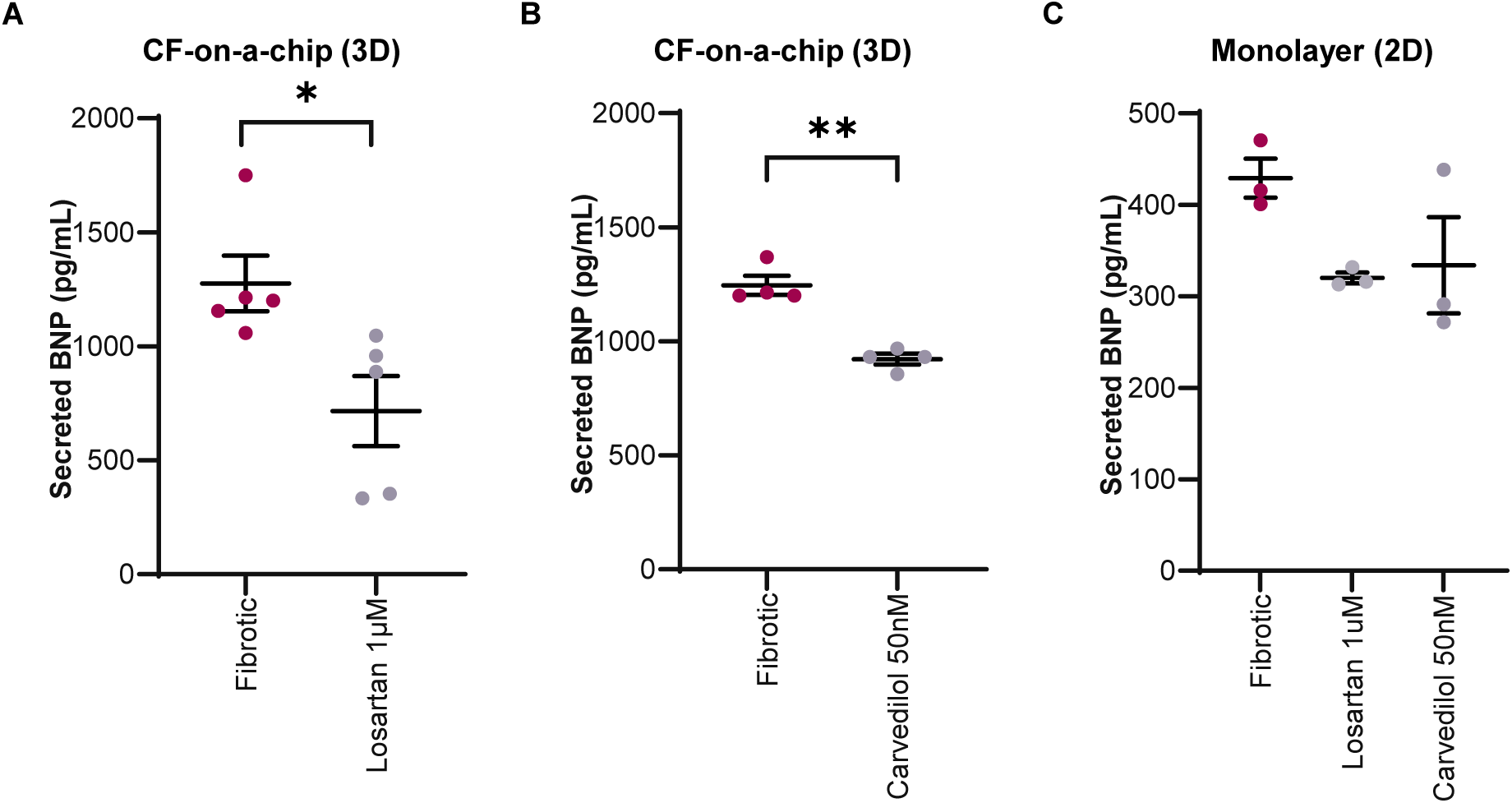
BNP secreted by losartan and carvedilol-treated fibrotic 2D cultures and 3D tissues (hCF-chip). **a**, BNP secreted by 1μM losartan-treated fibrotic tissues at day 21 vs fibrotic tissues (average ± SEM, p = 0.037, N = 5). **b**, BNP secreted by 50nM carvedilol-treated fibrotic tissues at day 21 vs fibrotic (average ± SEM, p = 0.0016, N = 4). **c**, BNP secreted by monolayer culture (1:3 human cardiac fibroblasts to hiPSC-CMs) for drug-treated fibrotic (TGF-β1-treated) monolayers vs. untreated fibrotic monolayers (average ± SEM, p=0.09 for losartan, p=0.17 for carvediolol; one way ANOVA, N = 3).

**Supplemental Figure 8.**
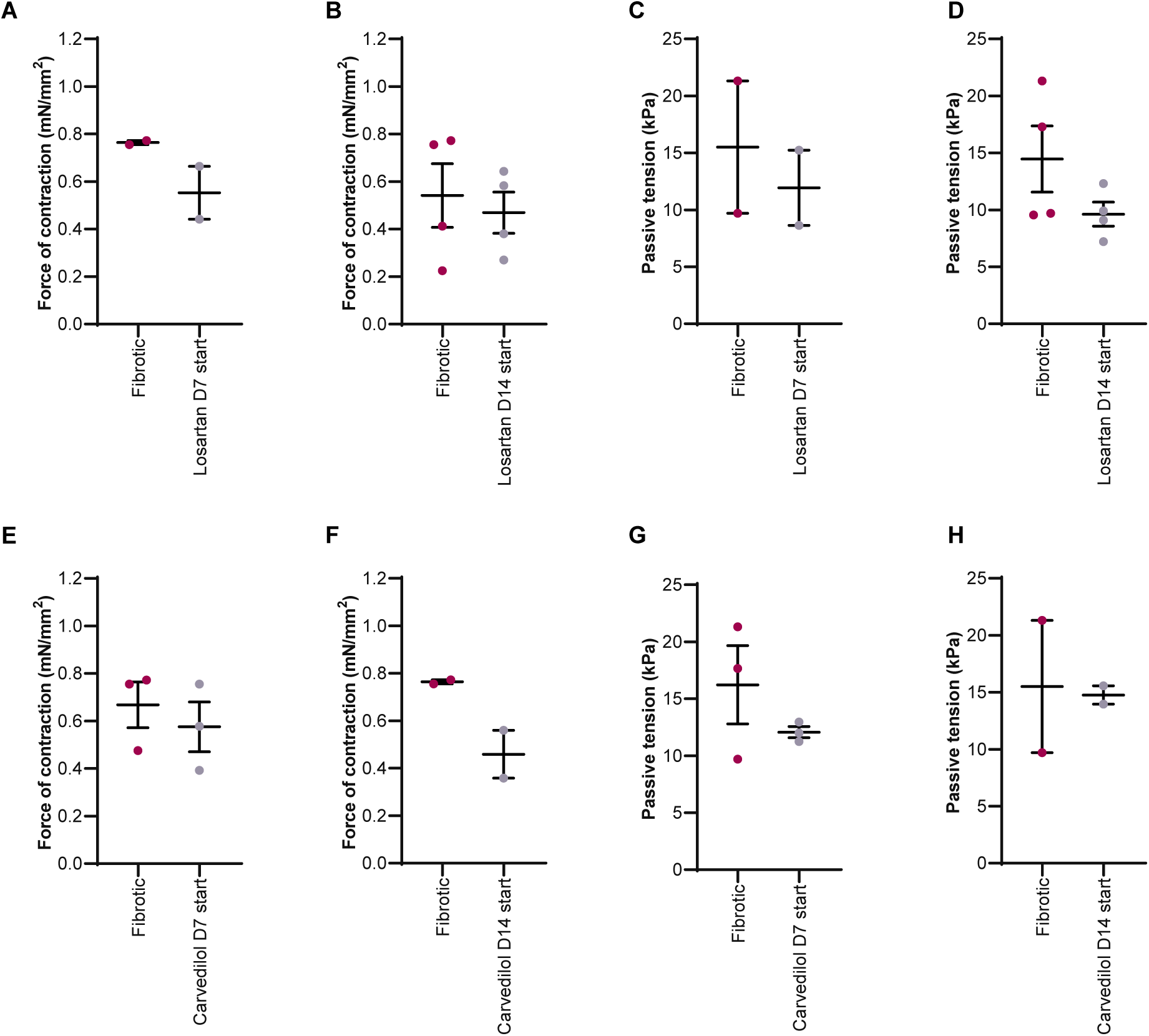
Treatment with losartan or carvediolol do not affect active force or passive tension. Assessment of active force and passive tension for losartan (**a** to **b**) and carvedilol (**e** to **h**) at day 21 post cell-seeding with treatments beginning from day 7 or day 14 post-cell seeding without TGF-β1 withdrawal (average ± SEM, N = 3 runs for **b, d, e**, and **g**, N = 2 for **a, c, f**, and **h**).

**Supplemental Figure 9.**
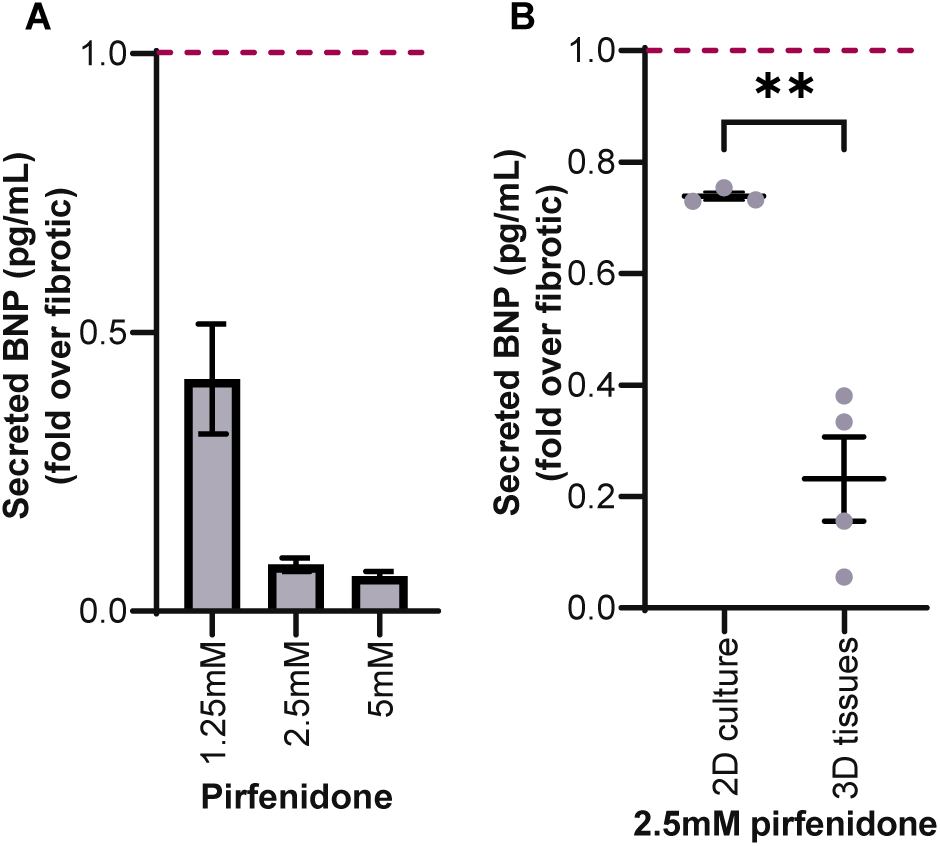
BNP secretion of pirfenidone treated hCF-on-a-chip tissues and 2D monolayers. **a**, BNP secretion at endpoint for 1-week drug treatment of fibrotic tissues with pirfenidone (N = 2), fold over untreated fibrotic (dashed red line) (average ± SEM). **b**, BNP secretion by fibrotic monolayers (TGFβ-treated) and fibrotic 3D tissues (hCF-chip) treated with 2.5mM pirfenidone, fold over fibrotic (dashed red line) (average ± SEM, p = 0.0024, N = 3 for 2D culture, N = 5 for 3D tissues).

**Supplemental Figure 10.**
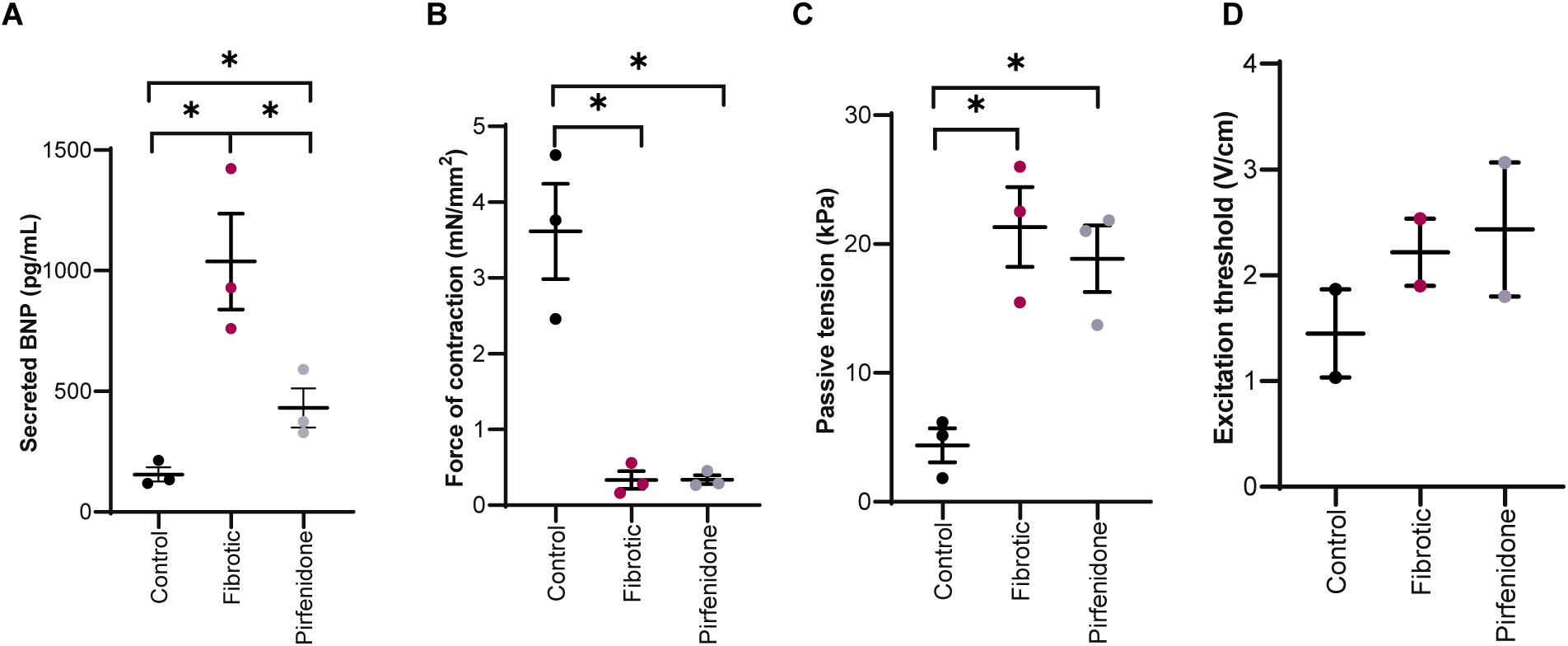
Evaluation of fibrotic response and drug treatment with an independent cell line (cell line B). **A**, BNP secretion (one-way ANOVA p = 0.034, t test p = 0.002 control vs fibrotic, p = 0.012 pirfenidone vs fibrotic) **b**, force of contraction (one-way ANOVA p = 0.043, t test p = 0.045 control vs fibrotic, N = 3), **c**, passive tension (one-way ANOVA p = 0.033, t test p = 0.036 control vs fibrotic N = 3), and **d**, excitation threshold (N = 2) show trends consistent with the main hiPSC-CM line used in the main figures (cell line A, Supplemental table 5); average ± SEM.

**Supplemental Figure 11.**
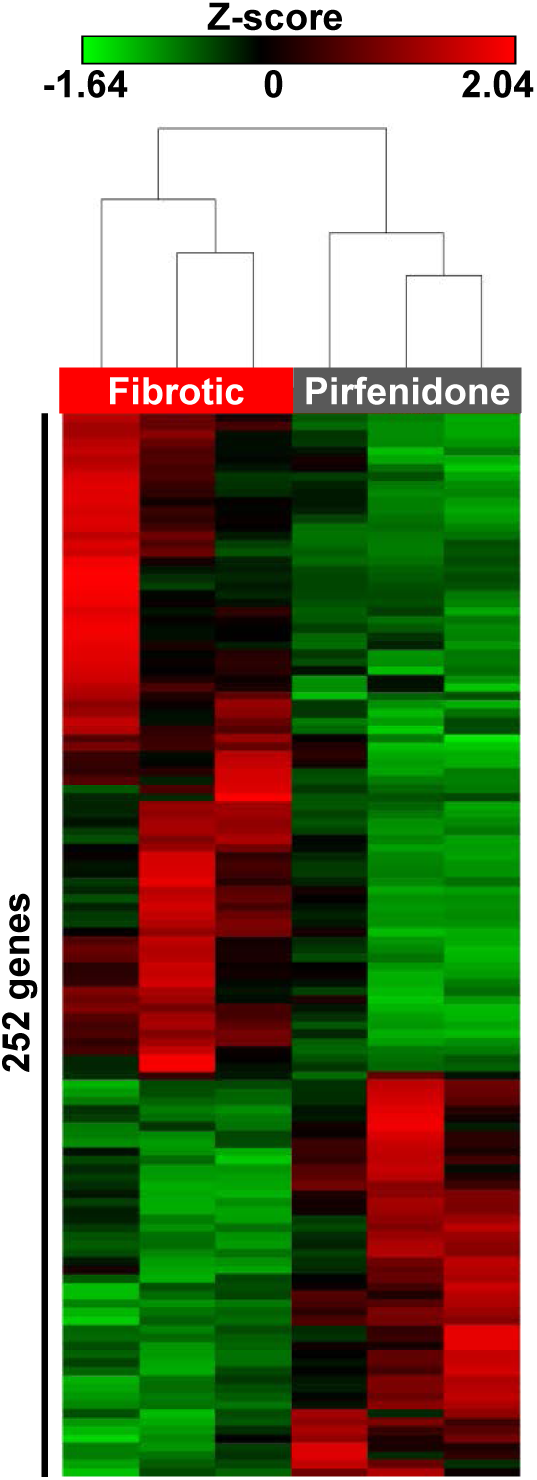
Transcriptomic analysis of control and fibrotic tissues by RNAseq. Unsupervised hierarchical clustering showing 252 differentially expressed genes in pirfenidone-treated and fibrotic tissues, z-scores for the RPKM (p≤0.05, ≥2-fold, N = 3).

**Supplemental Figure 12.**
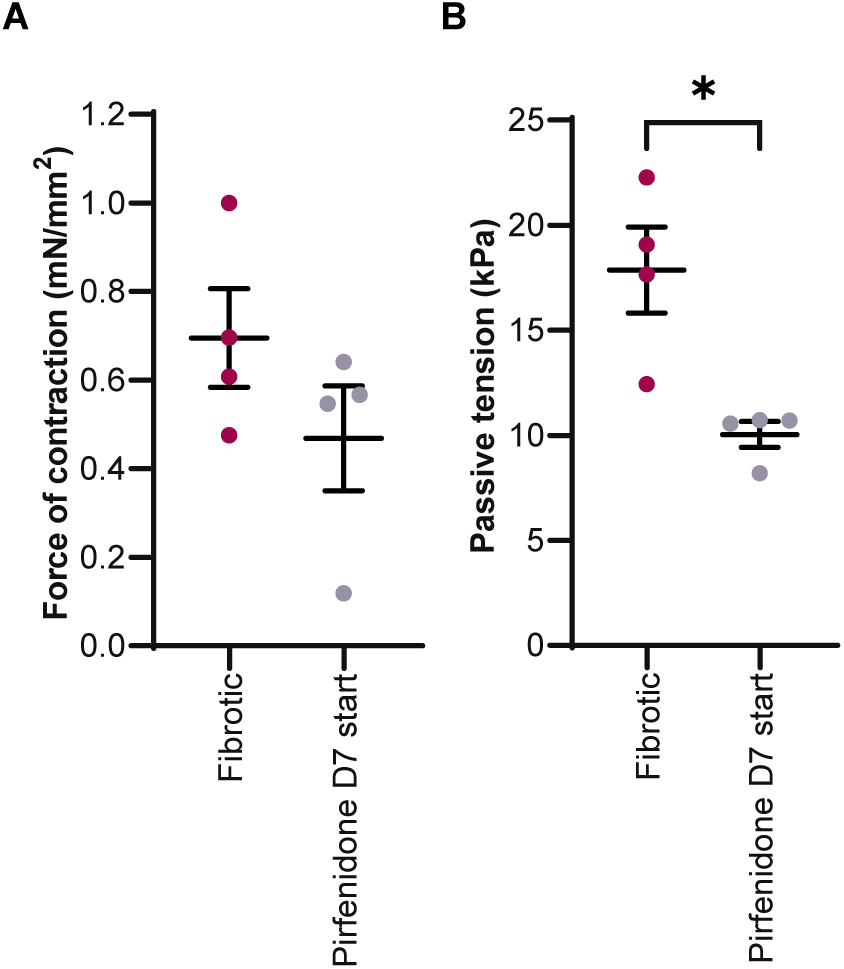
Treatment with pirfenidone for 2 weeks improves passive tension. Assessment of active force (**b**) and passive tension (**a**) for pirfenidone (2.5mM) with treatment beginning from day 7 post-cell seeding without TGF-β1 withdrawal (average ± SEM, not significant for force of contraction (left), p = 0.049 for passive tension (right), N = 4).

**Supplemental Figure 13.**
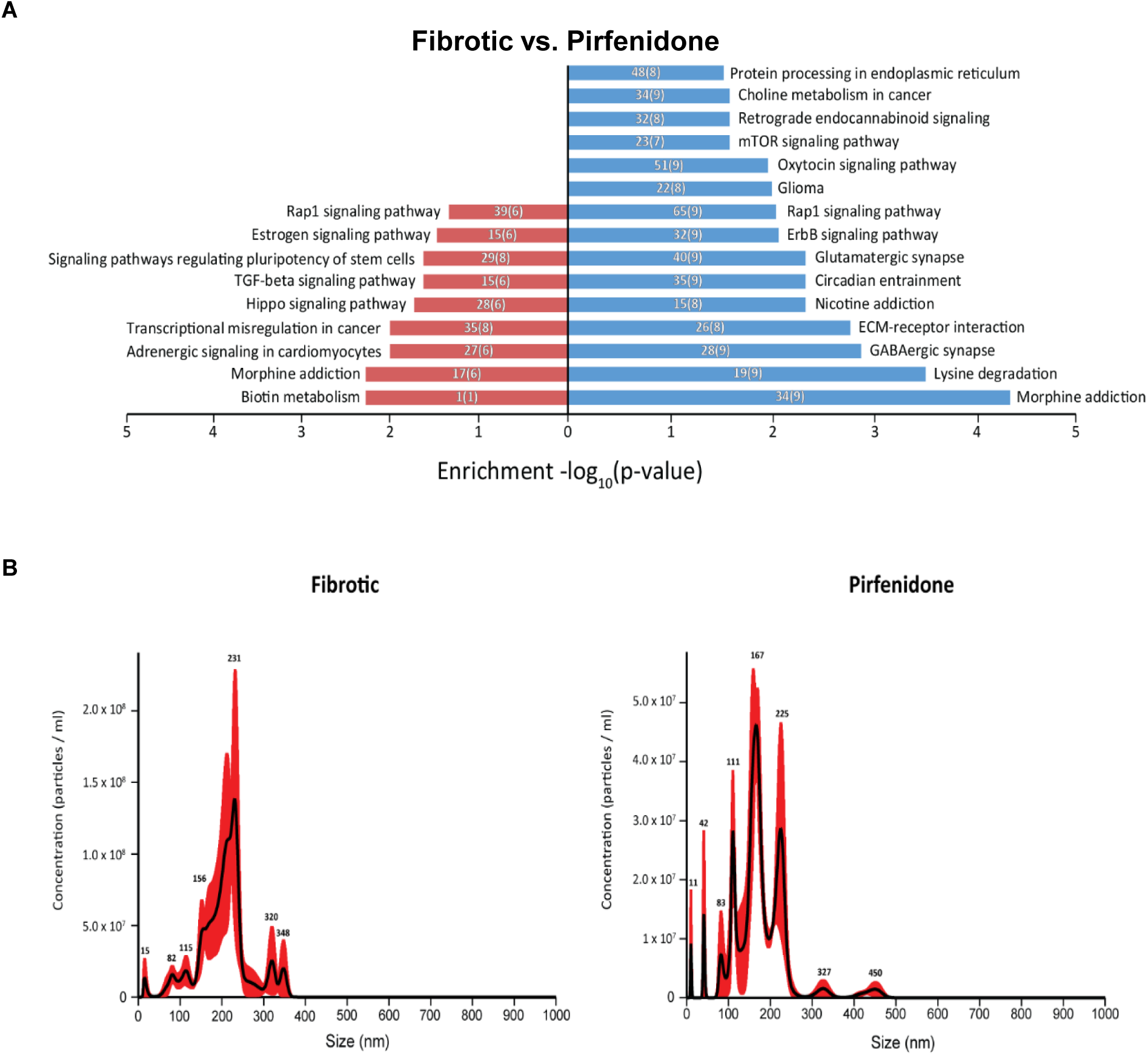
Pirfenidone-treated human CF-on-a-chip tissues exhibit a differential miRNA signature vs untreated fibrotic tissues. **a**, KEGG pathway analysis of signaling pathways predicted to be affected by microRNAs that were significantly down- (red) or up-(blue) regulated (N = 3). **b**, Representative size distribution of extracellular vesicles enriched from cell culture media from fibrotic and pirfenidone-treated tissues. The data were obtained from the binned averages of at least three 60 second NTA videos. The peak positions are marked. There are no statistical differences between groups for concentrations and 50 nm size bins. The black line represents the fitting curve of the binned data and the red error bars indicate ± SEM (N = 3).

